# Deep Learning enabled discovery of kinase drug targets in Pharos

**DOI:** 10.1101/2024.10.08.612754

**Authors:** Ádám M. Halász, Stephen L. Mathias, Srinjoy Das, Jeremy S. Edwards

## Abstract

We use machine learning with standardized molecular structure and gene ontology data to predict ligand interactions for a set of human kinases. We realize this by leveraging information from the TCRD / Pharos database, developed and maintained within the Illuminating the Druggable Genome (IDG) project.

Pharos collects relevant biochemical and clinically relevant information of a large set of biologically important (human) proteins from publicly available sources, including scientific publications as well as specialized databases. The 635 kinases listed in Pharos are classified into levels reflecting the relative amount and type of accumulated information. Importantly, molecular structure and Gene Ontology annotations are available for the entire set, but only 455 of the kinases have recorded ligand affinity data.

We developed a deep neural network-based framework to predict the ligand affinity profile for kinases using generally available information (molecular structure and Gene Ontology annotations) as input. The input data is organized into a 2,770 – dimensional vector with binary entries. The output data are predicted affinity values for interactions between the respective kinase and possible ligands.

To address the very large number of possible ligands (58,800) and the sparsity of available binding data, we organized the ligands into 5,275 clusters based on structural similarity measures. Our model framework is trained to predict likely interactions between kinases and these ligand clusters.

We aim to identify sets of likely ligand partners associated with high predicted relative affinities for a given kinase. We measure performance by evaluating the efficiency in identifying known ligand partners for documented kinases that were not included in the training data. Our results indicate that our model framework can identify sets of ligands that will contain a significant fraction of the correct (known) ligand partners.

## Introduction

The past few decades have seen the biological and biomedical sciences transform into a data rich discipline. Advances in bioinformatics, genomics, and DNA sequencing technology have revolutionized our ability to sequence and analyze genomes, edit genes, and manipulate individual proteins. Alongside these, significant increases in computing power and data storage capacities and algorithmic innovations have enabled the growth of data science, machine learning, and generative AI, pushing the boundaries of what can be achieved in these fields.

A particularly exciting frontier is the intersection of “omics” data—comprehensive collections of related biological data, like genomics, proteomics, and metabolomics—with machine learning and automated reasoning. This convergence is poised to fundamentally change our approach to understanding biological systems and enhancing drug discovery processes. Through these tools, we can connect complex datasets to predictive models that can suggest new hypotheses and accelerate the discovery of therapeutic targets and treatments (Hasin et al., 2017).

Pharos, as the user interface of the Knowledge Management Center (KMC) for the Illuminating the Druggable Genome project funded by the NIH Common Fund, serves as a key resource. Pharos is one modality to access the Target Central Resource Database (TCRD) which provides structured information about proteins, e.g. receptors, that are encoded by the human genome. Pharos/TCRD not only offers fundamental structural information, such as the amino acid sequences of these proteins, but also enriches this with gene ontology annotations and a range of biologically and clinically relevant data, from tissue expression patterns to ligand binding details (Kelleher et al., 2023) (Oprea et al., 2024).

For proteins, knowledge of chemical interaction partners (ligands) is a pre-requisite to uncovering their biological function and identification of modalities to modulate it. In drug discovery, identifying and understanding ligands that bind to these targets is crucial. Modern pharmaceuticals, such as the GLP-1 receptor agonists used in diabetes treatment, function by modulating the activity of specific (target) biomolecules. The development of such drugs follows the identification of therapeutic targets; for example, the GLP-1 receptor was identified as a target a decade before the emergence of drugs like semaglutide (Garber, 2011). The process of drug discovery involves narrowing down potential candidates through successive, resource-intensive steps, where having a chemical affinity of a given ligand to the target is a fundamental requirement (Mullard, 2020).

Ligand-target binding is fundamentally a physical interaction characterized by quantitative measures such as the binding energy; such measures can be estimated through molecular dynamics simulations (Tropsha et al., 2024) or by using machine learning based frameworks such as AlphaFold (Jumper et al., 2021), (Abramson et al., 2024). Our approach aims to complement these existing methods which use detailed protein sequence and genomic information, by relying on higher-level characterization of target and ligand molecular structure and biological function, that reflect accumulated domain-specific knowledge. By contrast with approaches involving detailed 3-dimensional structure, we use curated high-level structure information along with gene ontology annotations to predict likely ligand binding partners for a given molecular target. Our machine learning model is trained using published ligand binding data compiled and made publicly available in Pharos (Kelleher et al., 2023) (Oprea et al., 2024).

This paper explores the potential of Deep Learning to enhance the prediction of drug target interactions using Pharos and similar resources that compile and organize vast amounts of accumulated domain-specific knowledge. By integrating these powerful tools, we aim to help streamline the discovery process and highlight promising candidates more efficiently. While not meant to replace more traditional methods like QSAR, binding energy studies, and other approaches that rely on detailed structural and sequence data, our approach serves as a complementary strategy that narrows the search space and refines our ability to predict interactions in the vast landscape of potential drug targets (Jumper et al., 2021) (Volkov et al., 2022).

## Methods

### Source of Input and Training Data

The data in TCRD is compiled from multiple public repositories. Domain, structure, and functional site annotations are from the following three sources: Pfam (Mistry et al., 2021), <u>InterPro</u> (Paysan-Lafosse et al., 2023), and <u>PROSITE</u> (Sigrist et al., 2012). <u>Gene ontology</u> (GO) term annotations (Ashburner et al., 2000) (The Gene Ontology Consortium et al., 2023) for the kinases in TCRD were filtered as follows. Only leaf terms with experimental evidence codes were used, and only terms from the Biological Function and Molecular Process branches of the Gene Ontology were used.

Structural and gene ontology annotations for kinases were systematically extracted from the Target Central Resource Database (TCRD) version 6.13.4 using API queries. The extracted data are comprehensively summarized in two files: `TCRDv613_KinaseDomainAnnotations.tsv` for kinase domain annotations and `TCRDv613_GOAnnotations.tsv` and for gene ontology annotations. These files include detailed information on the structural attributes and biological functions associated with the kinases. Both of these files and the variables represented in the rows in columns can be obtained from our GitHub repository (https://github.com/amh65/KinaseMLPaper).

Ligand affinities for each target (kinase) and molecular structures specific to each corresponding ligand were obtained through structured queries to Pharos. The results of these queries are compiled in `LigandQuery.tsv`, which includes data on kinase-ligand interactions and identifying information as well as molecular characteristics of the respective ligands, and this file is available from our GitHub repository. This includes molecular structure encoded as SMILES (Weininger, 1988). The ligand molecular structures are analyzed using <u>RDKit</u> (Gori et al., 2022), a widely used open-source cheminformatics package. Molecular structures of the ligands are mapped as Morgan fingerprints (Morgan, 1965) and Tanimoto similarity (Bajusz et al., 2015) is used to cluster the ligands via Butina clustering (Butina, 1999).

The dataset is organized by kinase. Structural and gene ontology information is available for all 635 kinases included in the study. However, ligand binding information is available for only a subset of these kinases, specifically 455, which are referred to as the “documented” set.

### Model development

#### Data Processing for ML

The source data, which include structural information for kinases and ligands, gene ontology annotations for kinases, and kinase-ligand affinity measures, were processed and compiled into structured vector formats. This structured data representation facilitates the application of machine learning techniques for predictive modeling and analysis. Summary information on the molecular structure and biological function for each kinase is encapsulated in a 2770-dimensional binary vector containing the corresponding structural and gene ontology annotations. This representation was applied uniformly across all 635 kinases in the study. The kinase-ligand interactions for the subset of 455 kinases with documented ligand interactions are represented as a 5275-dimensional vector, containing representative affinities of the kinase to a group of similar ligands (see below for details on ligand groups). These processed datasets serve as the input for developing and training the machine learning models aimed at predicting potential drug targets and understanding the biochemical interactions at a detailed level.

##### Input Data Representation

The input data for the modeling framework is compiled in `TCRDv613_KinaseDomainAnnotations.tsv`. The file contains identifying details for the kinases along with structural information encoded as binary vectors. This binary encoding results in a comprehensive representation of kinase structural attributes.

From these data, we construct a 635×1091 binary matrix, denoted as `TargetVec`. This matrix encapsulates the structural information for all 635 kinases. Notably, `TargetVec` has a matrix rank of 383 and contains 399 unique rows, indicating the presence of redundant information among some kinases. For the subset of 455 kinases with known interactions^1^, the rank of the corresponding matrix reduces to 281, with 291 unique rows, highlighting a similar pattern of redundancy.

Additional features are obtained from gene ontology annotations in `TCRDv613_GOAnnotations.tsv`. In this file, each kinase’s gene ontology annotations are encoded in binary vectors, extending each kinase’s feature vector by 1,679 components. Thus, the original structural vectors are augmented to form a combined feature vector of 2,770 components. The resulting data structure is a 635×2770 matrix, referred to as `JVec`. This matrix combines the structural and gene ontology information into a single unified representation. `JVec` has 581 unique rows and a rank of 577, illustrating the high dimensionality and the richness of the dataset in capturing diverse biological phenomena. When this matrix is restricted to the 455 well-studied kinases, it has 426 unique rows and a rank of 425, maintaining a similar profile of uniqueness and redundancy as seen in the complete set of kinases.

##### Output Data Representation

The ligand interaction profiles were obtained through a direct query in the Pharos interface, with the results stored in the `LigandQuery.tsv` file. These profiles detail the chemical affinities between kinases and their ligands, characterized by five different measures (activity types): AC50, EC50, IC50, *K*_*d*_, and *K*_*i*_. Each of these measures corresponds to the concentration of the ligand at which 50% of a maximum response level is observed. For instance, the defining relationship for EC50 can be expressed as 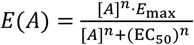 where *E, E*_*max*_ refer to the observed effect, [*A*] is the concentration of the ligand and *n* is the Hill coefficient (if any). The affinity measure is the negative 10-based log of the corresponding molarity^2^.

There are 106,518 individual kinase-ligand interactions involving the 455 interacting kinases and one of the 58,798 distinct ligands involved. When arranged by ligand and activity type, this results in a 455 × 239990 matrix with 1.33 × 10^8^ entries – one in 1255 of them nonzero. For our purpose – the identification of compounds that exhibit *any* type of chemical interaction with the targets – we merged the different types of activity measure by assigning the highest affinity of any type recorded for a given kinase-ligand combination. This results in the 455×58,798 activity matrix discussed below.

###### Dimensional reduction (1): Ligand Clusters / Groups

There are 58,798 kinase ligands referenced in the data initially extracted from Pharos. The affinity matrix, which incorporates all the activity types recorded, is very sparse. For the 455 kinases with any ligand activity information, the 455×58,798 matrix has 80,878 non-zero entries. This translates to an average of one non-zero entry for every 332 positions, or 0.31% non-zero entries. Additionally, the number of kinases (455 with ligand information) is relatively small in comparison to the dimension of the ligand affinity vectors.

To help address the dimensionality challenge, we used the ligand structural data to group ligands into clusters based on structural similarity. We used standard RDKit functions to obtain Morgan fingerprints (Morgan, 1965), compute pairwise Tanimoto similarity measures (Bajusz et al., 2015) and perform Butina clustering (Butina, 1999).

This resulted in 5,275 ligand clusters or groups. We assigned the highest recorded affinity between a kinase and any of the ligands in a group to represent the interaction between a kinase and a ligand group. The 455×5,275 matrix describing these groups has 13,211 non-zero entries, which corresponds to 0.55% occupancy. This reduction lowers the number of columns by approximately tenfold, though the matrix remains sparse with one in ∼180 entries being non-zero.

###### Dimensional Reduction (2): Projection

After assembling the combined structural / gene ontology annotation and the ligand-group affinity matrices, we are facing a high-dimensional prediction problem where the number of input and output dimensions, which are 2770 and 5275 respectively, are an order of magnitude higher than the available number of samples which is 455.

To further reduce the dimensionality of the feature and output vectors, we project them onto chosen basis sets (Shlens, 2014). The standard approach for this is to use Singular Value Decomposition (SVD), most often used in statistics as part of Principal Component Analysis (PCA) where the vectors are first shifted by subtracting their mean over the entire group. MATLAB’s built-in PCA function returns the complete set of principal vectors and corresponding loadings for a set of vectors, 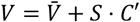. Here *V*[*N*_*K*_ × *N*_*LG*_] contains the original vectors as rows; *C*[*N*_*LG*_ × *N*_*D*_ ] contains the basis vectors 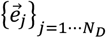 as columns (when using PCA, these are the same as the principal vectors); *S*[*N*_*K*_ × *N*_*D*_] contains the coordinate vectors (also in columns); finally, 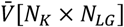 contains the mean ligand vector 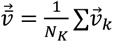 in each of its rows. A given vector 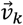 is represented as

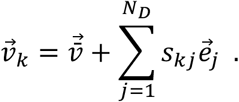

Given the limitations of our machine learning framework, training a neural network using all *≈* 350 − 400 components of the input and output coordinate vectors poses a technical challenge. A feasible workaround is to use only a subset of the respective components, truncating the representation to typically 100 – 200 components on the input side and 20 – 200 on the output side.

#### Neural Net Model

The input and output vectors are provided in the csv files `NNInputVec.csv` and `NNOutputVec.csv`. Input vectors are available for all 635 kinases, but the output vectors (which reflect documented interaction affinities with one of the 5275 ligand groups) are only available for the 455 documented kinases. The SVD basis on the output side is derived from the PCA decomposition of the training set only and it is dependent on the choice of this set.

##### Truncated SVD projections

Our objective was to develop machine learning model 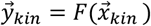 scapable of predicting the ligand binding affinity vector 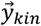, for a specific kinase, from the molecular structure / gene ontology vector 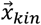 associated with that kinase. As mentioned earlier, we utilized truncated Singular Value Decomposition (SVD) to project both the kinase structure and ligand group affinity vectors onto a reduced-dimensional space, typically involving a subset of their respective singular vectors.

The input vectors, representing kinase structure and gene ontology, are inherently binary, with components taking values in {0,1}.

To standardize the analysis and enhance the interpretability of the model’s predictions, we first rescaled the original ligand affinity vectors to the range [0,1]. This normalization was performed component-wise, ensuring that the highest recorded affinity for each specific ligand group across all kinases was scaled to exactly 1. Importantly, this scaling process preserved zero affinity values, maintaining the pattern of sparsity in the dataset. The quasi-binary nature of the affinity vectors simplifies the modeling process and provides a clear, interpretable baseline from which deviations due to affinity can be measured.

We use these scaled representations as a reference when comparing predicted affinities to known experimental values. This approach ensures consistency and aids in the qualitative and quantitative assessment of our machine learning models’ performance in predicting ligand binding affinities.

We performed **PCA based projections** as described in the previous section to obtain lower dimensional coordinate vectors 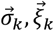 to represent the input and output vectors with respect to the bases 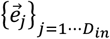 and 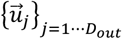 respectively.

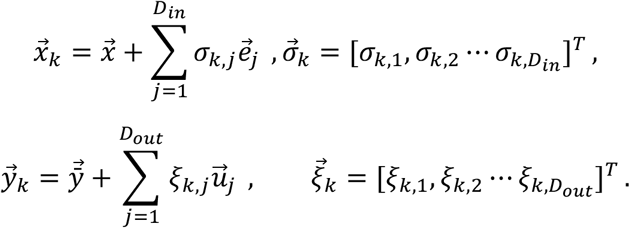

For the set of 455 documented kinases, the full dimension of the input and output coordinate vectors is *D*_*in*_ = 425 respectively *D*_*out*_ = 289. ^3^

Finally, the **coordinate vectors** 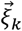 representing ligand (group) affinities are **rescaled** into the [0,1] interval (component-wise across all kinases, so that the minimum and maximum coefficient for each (principal) component is 0,1 respectively). This step, commonly used in statistical modeling (Goodfellow et al., 2016) improves the training and performance of the neural net model.

We trained neural net models^4^ using **truncated coordinate vectors**, retaining only a subset of the SVD coordinates, of dimensions typically *N*_1_ = 100 … 200 respectively *N*_2_ = 20 … 289. We refer to these as input / output sizes.

#### Multilayer Neural Network Architecture

##### Core Model Structure

The foundational structure of our model is a feed-forward neural network with 2 - 4 hidden layers, in addition to the input and output layers of sizes, *N*_1_, *N*_2_. A representative neural network configuration size is {100, 1000, 500, 200, 20} (with *N*_1_ = 100, *N*_2_ = 20). The dataset comprising 455 vector pairs, which correspond to the documented kinases, is divided into training, validation, and test sets with a ratio of 70:15:15. The optimization of the network parameters is performed using scaled conjugate gradient descent.

##### Model Configuration Exploration

We conducted experiments with various network configurations and dimensions of the input/output vectors. These experiments included multiple runs with consistent partitioning of the data into training, validation, and test sets to ensure the reliability and reproducibility of our results.

##### Training and Validation Process

A critical aspect of our training and validation process is designed to simulate the real-world application of the model, particularly for predicting binding partners for kinases that lack documented ligand affinities. Therefore, for the kinases in the test set, we treat their interaction vectors as entirely unknown; these vectors are not included in the Principal Component Analysis (PCA) basis used for dimensionality reduction.

Moreover, any ligands that interact exclusively with the test-set kinases (and not with those in the training or validation sets) are omitted from our analysis.

This exclusion is specific to the kinase affinity vectors pertaining to the test set. For the intended application, the validation set must include known kinases, hence their ligand interactions are incorporated into the model. However, the input vectors for all kinases, including those without documented ligand interactions, are known and used in the model.

#### Linear Regression Model

We developed a linear regression-based (LR) model that can work concurrently with the NN model described above. This model is trained on the same coordinate vectors 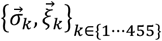 as the NN model. For a given instance of the LR model the sample vectors are truncated to retain only the first *N*_1_ ≤ *D*_*in*_, *N*_2_ ≤ *D*_*out*_ components respectively, similarly to the NN models. Unlike the NN model, the linear model is *trained separately for each output component j* = 1 … *N*_2_ using the complete sample inputs 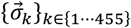 as predictors^5^ for the corresponding sample outputs of the respective component ={ξ_*k,j*_}_*k*∈{1…455}_. The resulting bias *b*_*j*_ and linear coefficient vector 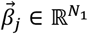 provide the LR prediction for coordinate *j* of sample *k*:

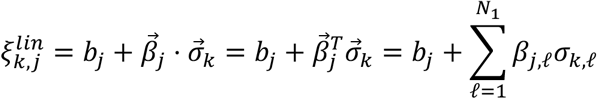

The bias and coefficient vectors (in row form 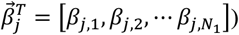 are conveniently organized into a bias vector and coefficient matrix with row index *j* = 1 … *N*_2_

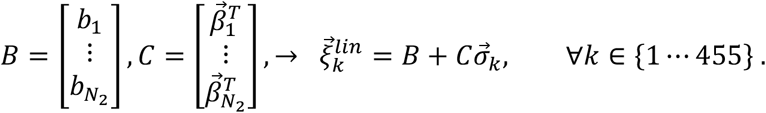

Each instance of the LR model is defined by these two objects 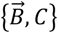 that are the combination of *N*_2_ <u>separately</u> trained linear regression models.

#### Reconstruction and Evaluation of Predicted Affinities

##### Reconstruction

Once trained, the Multilayer Perceptron (MLP) model predicts scaled coordinate vectors from the input vectors (or rather, from the input coordinate vectors) 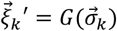. The output of the model is then adjusted to reverse the initial scaling and projection steps. This involves rescaling in the coordinate vector space and projecting back to the affinity space,

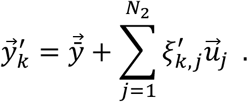

The resultant *predicted scaled affinity vectors* are directly comparable to those used during training.

##### Combining NN and LR models

The reconstruction and evaluation procedure for linear regression is identical, using 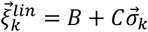 as the predicted (truncated) coordinate vectors.

We also generated *combined* NN / LR predictions by combining the predicted scaled affinity vectors 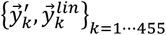 obtained from two selected models. For each component of each kinases’ affinity vector (*y*_*k,g*_ – scaled affinity of kinase *k* with respect to ligand group *g*), the *combined predicted affinity* is defined as the higher of the two model predictions:

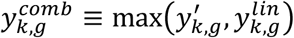

This procedure can be used for any two models. We analyzed combinations of pairs of NN and LR models on two configurations, (i.) one or both of the input and output dimensions of the LR model matched the sizes of the respective layers of the NN model and (ii.) combining each NN model with the best performing (typically largest size) LR model.

##### Evaluation of Model Performance

The utility of the model is determined by its ability to identify ligand groups that interact with a kinase.

We compare a set of model-predicted affinity values 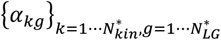 where *α*_*kg*_ refers to kinase *k* and ligand group *g* (the predicted affinity vector is 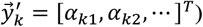, to the known kinase-ligand group pairs that have a recorded affinity 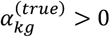.

For this purpose, we define a binary flag *δ*_*kg*_ = 1 if 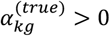 and *δ*_*kg*_ = 0 if 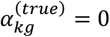. We refer to a group *g* as a “hit” for kinase *k* if *δ*_*kg*_ = 1, and a “nonhit” otherwise.

We use the predicted affinity vectors to infer “hits” based on the predicted affinity values. This setup forms a classic binary classification challenge, where the task is to discern positives and negatives based on a quantitative measure.

To quantitatively assess the performance of our model, we computed Receiver Operating Characteristic (ROC) curves (Hanley & McNeil, 1982) (James et al., 2023) for individual kinases and collectively for the entire dataset encompassing the training, validation, and test sets. Additionally, we calculated precision *TP*(*t*)/ (*TP*(*t*) + *FP*(*t*)) and recall *TP*(*t*) / (*TP*(*t*) + *FN*(*t*)) metrics^6^ for specific threshold values.

###### ROC curves

Given a set of predicted affinities {*α*_*kg*_} and a cutoff threshold value *t*, the putative “hits” or positives are those with *α*_*kg*_ > 0, and the rest are negatives or “nonhits”. Comparing this with the true labels *δ*_*kg*_, we have a number of true and false positives and negatives, that vary as the cutoff changes. The number of true positives as a fraction of the total number of hits is (also) called the true positive rate 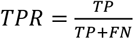; the number of false positives as a fraction of all negatives is called the false positive rate 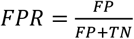. The ROC curve is a parametric curve in the (*FPR, TPR*) plane (false positives on the x-axis, true positives on the y-axis) obtained as the cutoff value *t* ranges from the smallest predicted affinity (*t* ≥ *t*_*min*_ = m*in*({*α*_*kg*_})) to the highest one (*t* ≤ *t*_*max*_= *max*({*α*_*kg*_})).^7^ The area under the ROC curve is a measure of the performance of the classification, with a value of 1 corresponding to perfect classification and 0.5 to random choice.

The true / false positive / negative classification can be performed for each kinase separately (considering the predicted affinities for fixed *k* and all relevant *g* indices) or for multiple kinases; the latter is most relevant when evaluating the performance over the training and test sets.

###### Precision and Recall with Percentage Cutoffs

In addition to the ROC curves, we computed precision and recall metrics as indicated above. Unlike the area under the ROC curve which is cutoff-independent, these measures are specific to a cutoff value. Since the range of the predicted affinity values varies significantly between individual kinases^8^, we used *relative* cutoff values at a fixed fraction (percentage) of the set of affinities being classified. We calculated precision and recall values corresponding to the top 5% and 1% of all predicted affinities (for individual kinases or for a group of kinases).

Consistent with the intended application of predicting as of yet unknown ligand partners to “dark” kinases, the corresponding cutoff values are accessible even without the knowledge of the actual hits.

For performance evaluation, we also computed precision values (*TP*/(*TP* + *FP*) for a cutoff *t*^∗^ that yields half of the known hits (i.e. 50% recall). Obviously, this cutoff value is not available for unknown kinases, but the measure is useful to compare performance for kinases whose number of known hits ranges from 1-3 into the hundreds. With approximately 5000 interacting ligands, 5% corresponds to 250 hits and 1% to 50 hits. Thus, a kinase with only 5 hits would have a precision measure of ≤ 0.02 or ≤ 0.10 respectively, even if the recall is 100%.

## Results

Our study primarily focuses on human protein kinases, which are enzymes that catalyze the phosphorylation of other molecules, playing a crucial role in various cellular processes. Within the Pharos database, there are a total of 635 entries classified under the kinase family of targets. These entries correspond to the known protein kinases encoded by the human genome.

The kinases are systematically categorized based on the extent of experimental and functional information available for each kinase, often referred to as their “illumination” level (Oprea et al., 2024). Specifically, they are distributed as follows; **Tclin -** 68 kinases with extensive clinical and functional data, **Tchem -** 387 kinases with significant chemical information primarily related to ligand interaction, **Tbio -** 161 kinases with biological data but lacking comprehensive ligand binding information, **Tdark -** 19 kinases with minimal available information and no recorded ligand binding data.

It is noteworthy that ligand binding information is not recorded for kinases categorized under Tbio and Tdark, highlighting a gap in the data for these lesser-known kinases.

### Ligand Binding Information and Analysis

#### Overview of Ligand Binding Affinities

Ligand binding information is quantitatively described using log-affinity values, which are compiled from a range of published research articles. Although multiple activity types are reported, all affinities are presented as numerical values. These values correspond to the ligand concentration that achieves 50% of the maximum effect on the kinase, measured in molar units. Due to the typically small concentration values, ranging from micromolar to nanomolar (even picomolar) levels, affinities are expressed as the negative of the base-10 logarithm of the molar concentration, *A* = −*log*_10_(*C*_50_[*M*]). While there are multiple activity (assay) types recorded in TCRD / Pharos, and in some instances, multiple entries for the same interaction type and target-ligand pair, we assigned the highest affinity of all types and instances for a given interacting pair.

#### Distribution and Policy for Including Affinity Values

The number of recorded ligand partners for each kinase varies widely, with 133 kinases interacting with only 1 to 5 ligands, while others, like JAK1, have as many as 3,142 interactions. This is significantly fewer than the total number of referenced ligands, which stands at 58,798. The distribution of affinity values notably declines below *A* = 7.3 reflecting a kinase-specific cutoff of [*C*50] ≥ 30*n*M with the exception of approved drugs with a well-understood mechanism of action (Oprea et al., 2024).

Consequently, the absence of a recorded affinity between a specific kinase-ligand pair can indicate a lack of available information or may reflect experimental investigation[s] that resulted in a low affinity value that was either unpublished or excluded from Pharos. In our analyses, we must keep in mind that we implicitly assigned an affinity value of 0 to kinase-ligand pairs for which no data is available.

#### Sparse Nature of the Kinase-Ligand Affinity Matrix

The kinase-ligand affinity matrix used as our starting point is sparse, with only 80,878 nonzero entries in a 455 × 58798 matrix. This results in an occupancy of just 0.302%. The rank of this matrix is 419, indicating that the majority of kinases have unique activity profiles. However, with an average of 1.375 kinase interactions per ligand, it is evident that subsets of the 455 documented kinases, such as those used for training and validation will have a significant number of non-interacting ligands.

### Ligand Clusters / Groups

#### Dimensionality Reduction through Ligand Clustering

To manage the high dimensionality of the ligand affinity vectors, we utilized ligand clusters based on the ligands’ structural similarities. Molecular structure information is accessible for all referenced ligands, and we employed Tanimoto similarity (Bajusz et al., 2015) to group ligands into clusters using Butina clustering (Butina, 1999). This approach resulted in a total of 5,275 ligand clusters (or groups).

For each kinase and ligand cluster, we assigned a representative affinity value. This was determined by identifying the highest recorded affinity for that kinase with any of the ligands within the cluster. The subsequent modeling and analysis are based on these derived ligand group affinity vectors.

#### Variability in the Number of Recorded Ligand-Kinase Interactions

The range of recorded ligand interactions for a single kinase varies significantly, from as few as 1 – 3 interactions to several thousand. For instance, JAK1 has 3,142 interactions, JAK2 has 2,858, PIM1 has 2,741, VGFR2 has 2,707, and EGFR has 2,527. On average, the 455 documented kinases each have 183.7 ligand interactions, with a median of 18. This variability probably reflects the extent of research devoted to studying specific kinases, rather than an intrinsic propensity of the kinases for ligand binding.

#### Distribution of Ligand Partners Among Kinases

The distributions in Figure S1 highlight a pronounced imbalance in the distribution of known ligand partners among kinases. Histograms of the number of kinases by ligand count ranges show that the majority of kinases have fewer than 100 interactions, with a median of 18 interactions per kinase. However, the average is 182.77 ligand interactions per kinase, indicating that the majority of known interactions are concentrated in a small subset of kinases.

The kinase-by-kinase distribution of interactions (Figure S2, with kinases sorted by the number of known interactions) shows a sharp decline in the number of interactions beyond the first 50-100 kinases. Cumulative distributions reveal that a minority (64 and 120, respectively) of the 455 documented kinases account for 80% of all interactions. It is crucial to keep this imbalance in mind when constructing and interpreting predictive models for kinase-ligand interactions.

#### Ligand Clusters Reflect Distribution Features

The plots in Figures S1 and S2 show the corresponding distributions of kinase interaction counts using individual ligands and ligand clusters. The skewed distribution of interactions observed with individual ligands is mirrored in the plots for ligand clusters, though the imbalance is somewhat reduced. The number of kinases responsible for 80% of all interactions nearly doubles when using ligand groups, illustrating that ligand clustering can help mitigate some of the distributional skewness.

#### Predicted Affinity Vectors

We developed a deep learning-based model to predict ligand group binding affinities from kinase structure and gene ontology annotation vectors. The dataset comprises 455 pairs of input and output vectors, which we split into training, validation, and test sets. We project both the input and output sets of vectors onto a basis spanned by their^10^ principal components (obtained through PCA). The coordinate vectors are then truncated, keeping only a smaller number of components, and these truncated vectors are used to train a multi-layer feed-forward neural network (multilayer perceptron or MLP).

#### Known Interaction Partners Can Be Recovered from Model-Predicted Affinities

We present illustrative results from a neural net configuration^11^ that uses 100 principal components for input and 40 for output, with three hidden layers consisting of 1000, 500, and 200 neurons, with randomly selected train (319 kinases), validation and test sets (68 kinases each).

The primary items of interest are the predicted ligand affinities, specifically the affinities between a kinase and a ligand group (LG). It is important to note that the original affinity values, which range from 4 to 12, are transformed into scaled affinity values (SA), such that the highest affinity for a given LG is normalized to 1. Non-interacting ligand groups correspond to an affinity of zero, for both scaled and unscaled values.

Ligand interactions that have been measured are included in TCRD / Pharos if the corresponding affinities are above a certain cutoff value, thus the *a priori* classification of kinase – ligand group pairs as interacting or non-interacting is based on an underlying continuous quantity, the true affinity (where it is known).

Our objective is to identify potential, yet unknown, binding partners for kinases. The performance and utility of the model are assessed by comparing the predicted interactions with the known interactions for the respective kinases. This comparison is most informative for kinases that were excluded from the model’s construction and training phases (i.e., the test set), but we also examine results for kinases in the training and validation sets.

As depicted in Figure 3, model-predicted affinities (for a given kinase) are generally higher for interacting ligand groups (LGs), although non-interacting LGs typically exhibit nonzero predicted affinities. The effectiveness of the model lies in its ability to use the predicted SA values to identify interaction partners, i.e., recover or predict ligand clusters that demonstrate significant affinity for the kinases of interest.

**Fig. 1:**
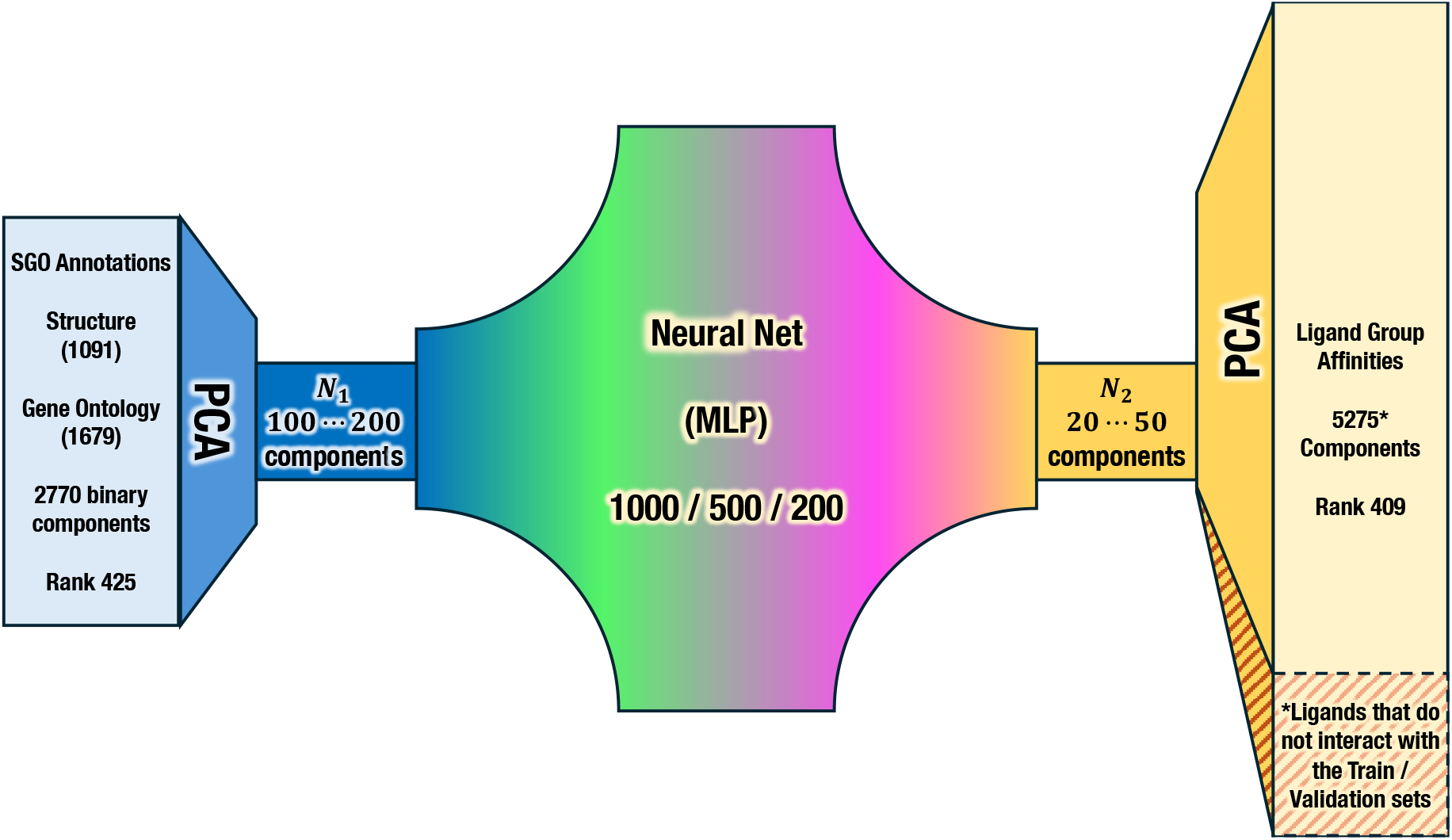
Schematic of the multilayer perceptron-based model outlining the mapping 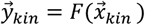 from the 2770-dimensional input (structural + gene ontology annotation vectors) to the 5275-dimensional output (ligand group affinity) vectors. Both sets are projected onto a basis of their span* (obtained through PCA). The resulting coordinate vectors are truncated and only the first N_1_, respectively N_2_ components are used in the NN model. *The test set affinity ligand group affinity vectors are not included when constructing the projection (PCA) basis; ligands that only interact with the test set (with identically zero entries on the train and validation vectors) are excluded; the clipped test-set vectors are otherwise processed identically to the others.

**Figure 2:**
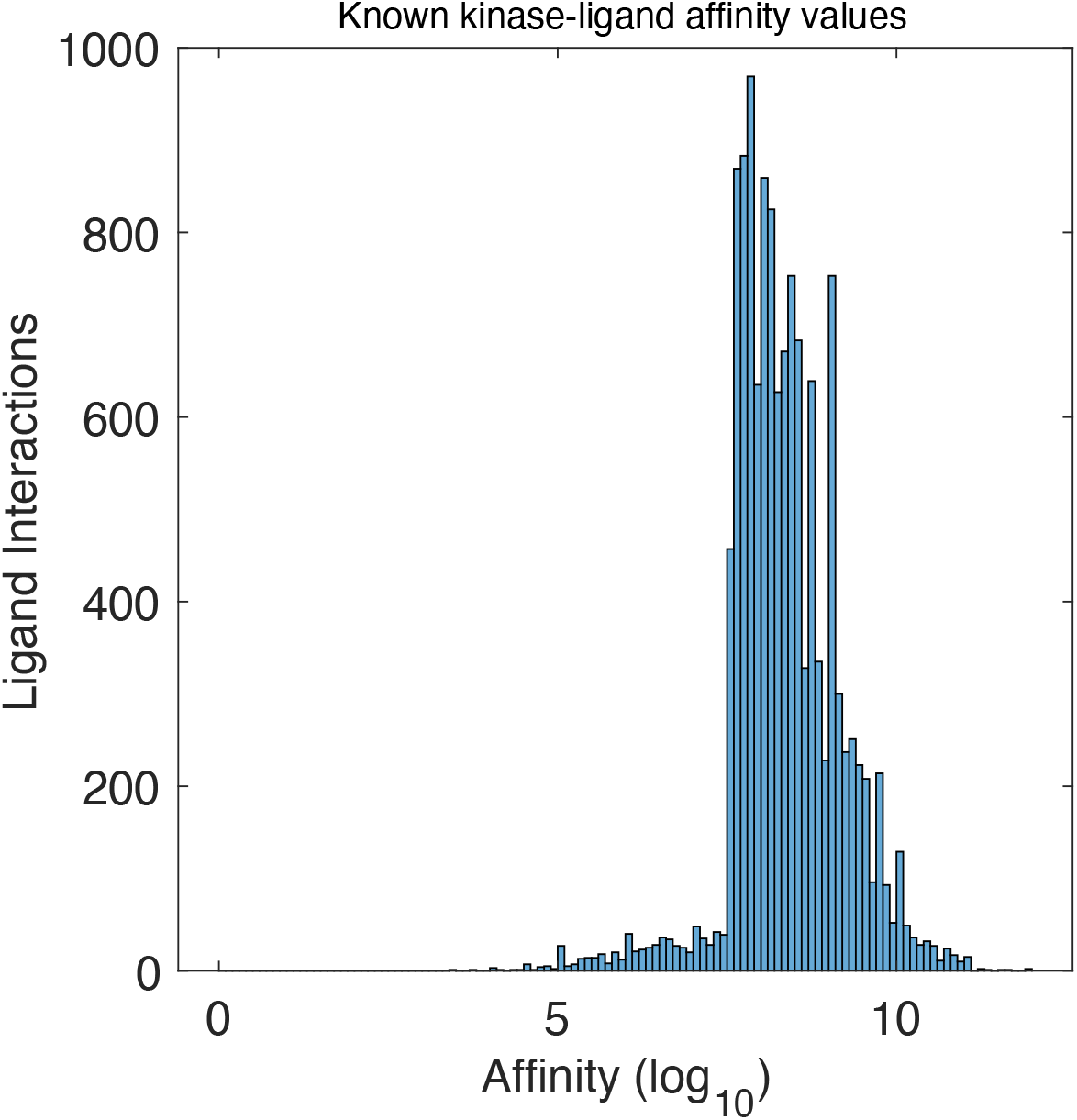
Distribution of the 80878 recorded (maximum) affinities of kinases to individual ligands. The shape of the distribution shows a significant drop below 7.3, consistent with the policy followed in compiling Pharos / TCRD data. It is also likely that many measured low affinity values were not published.

**Fig. 3:**
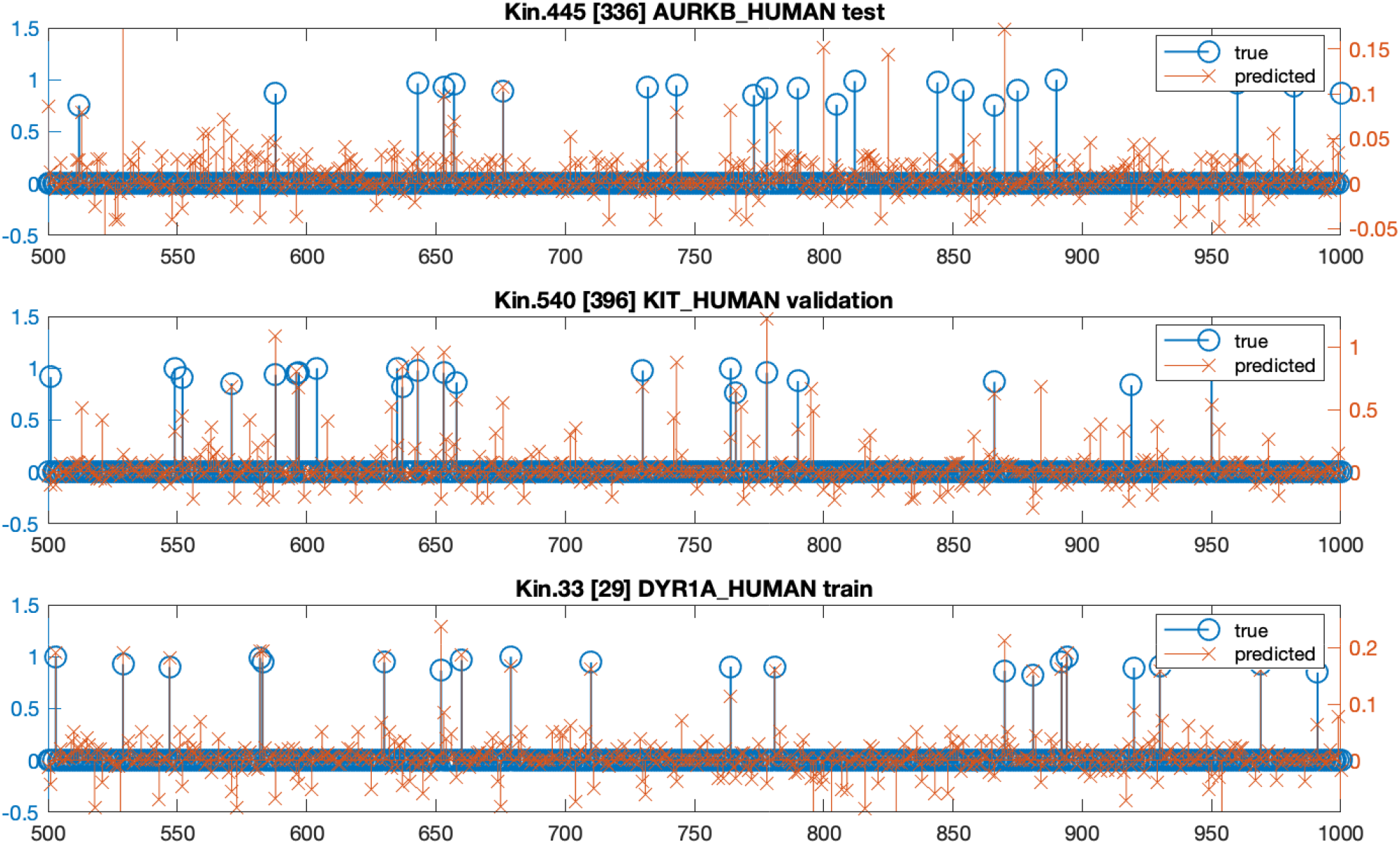
True and model predicted ligand group affinities for three kinases. The true affinities (left y-scale) used as training input are scaled to the [0,1] interval so that ligands with known interactions have nonzero affinities approaching 1; affinity values for ligands with no recorded interaction are exactly 0. The neural-net predicted affinities (y-scale on right-hand side) approximate the true values. For a given kinase, the predicted affinities induce a ranking of the ligand groups used to infer likely interactions. This is the basis of the performance measures discussed in the paper. Panels show different kinases; their role in the model training process is as indicated. This plot represents results from run #74839.^9^

**Fig. 4:**
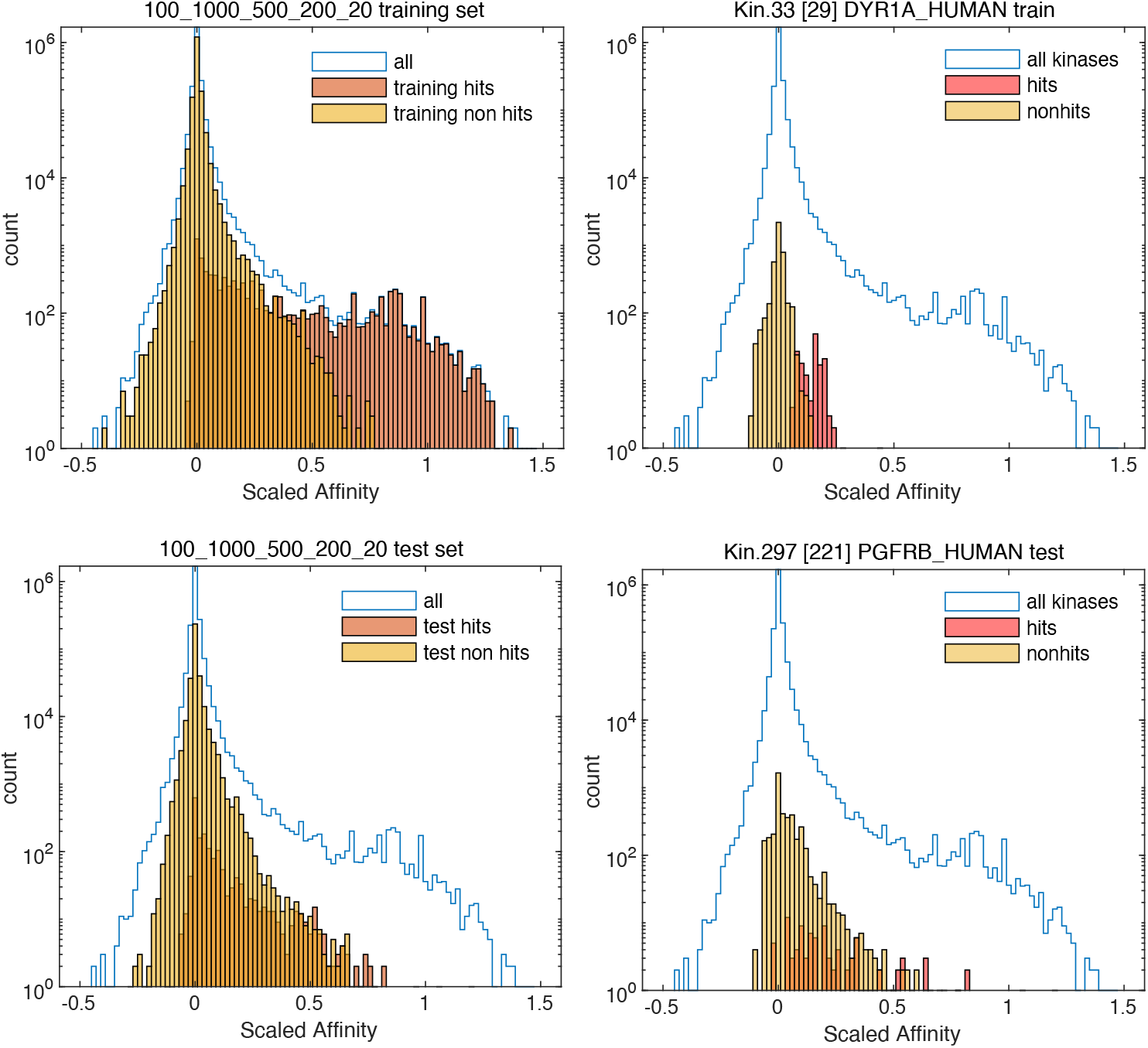
Distribution of model predicted LG affinities for the entire train and test set of kinases (left panel) and for one kinase from each group (right panel). Hits and non-hits are shown separately. The bulk of the distribution of predicted affinities for hits is higher than that of non-hits (note the logarithmic scale on the y axis), but many hits have predicted affinities overlapping with the bulk of non-hits. A clean separation only occurs in a subset of cases. We frame the identification of hits based on predicted affinities as a classic binary selection problem, whose outcome depends on the choice of a cutoff.

For a given kinase and its affinity vectors, we define a ligand group as a “hit” if there is a nonzero recorded (true) affinity^12^. The predicted affinities are then utilized to infer the “hits” for a kinase. We assess the performance of the model based on our ability to correctly identify these hits, i.e., to accurately determine the known interaction partners for kinases not used in the training phase.

The key factor that determines the extent to which model predicted LG affinities can be used to identify “hits” (interacting kinase – ligand group pairs) is whether the distribution of predicted affinities corresponding to hits is distinguishable from the rest (“non-hits”). Examples of such distributions for a single kinase and for the entire set are shown in Fig.4. For the <u>known affinities</u>, hits and nonhits have non-overlapping distributions – hits are in an interval that ranges from 1 down but never reaches zero, and nonhits have affinity exactly zero.

The model prediction maps the two subsets to qualitatively different distributions. The mass of the hit distribution is located higher on the affinity scale than that of the non-hits, but the two distributions overlap. The *nonhit distribution* is a single mode centered around zero and decreases on both sides somewhat slower than an exponential. The *hit distribution* has a narrow, peaked component similar to the non-hit distribution, but the bulk exhibits a slow decrease extending from 0 to a wide mode close to 1. Note that the vertical scale in Fig.4 is logarithmic. In the [0,0.3] range, the density of non-hits overwhelms the hits by more than 2 orders of magnitude. By contrast, for [0.5,1] and above, the hits are the majority, exceeding the density of nonhits by a factor of 10 or more from 0.7 and up.

#### Coordinate Vector Truncation vs. Neural Net Mapping Loss

It is worth taking a closer look at the sources of error impacting the predicted affinities. The original ligand group affinity vectors are scaled then shifted by a constant vector and projected onto a set of basis vectors as described previously^13^. The basis is obtained from the training and validation sets and the decomposition scheme spans those vectors – thus the representation:

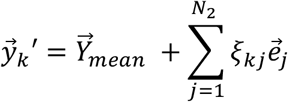

is loss-free 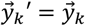 when using the complete set of basis (PCA / SVD) vectors, i.e. *N*_2_ matches the rank of the set of vectors used to construct the basis. For the 319 train vectors, this is *N*_B_ = 289. When *N*_2_ < *N*_B_, the reconstruction will be incomplete and the recovered vectors 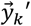 will be approximations of the true 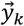.

The neural net (MLP) is trained to predict the *truncated coordinate vectors* 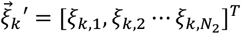 from the (also truncated) input coordinate vectors 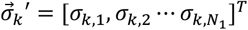. Thus, a perfectly performing MLP would reproduce the truncated coordinate vectors 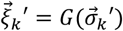 which would then map to the approximate scaled affinity vectors 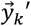. The real-world model will provide an approximation 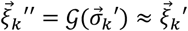 which in turn maps to the output 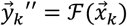.

It is useful to compare the exact 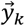 and predicted 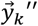 affinity vectors with the intermediate 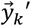 for specific kinases with attention to affinities corresponding to hit / non-hit ligands. This is illustrated in Fig.S3.

The kinase DYR1A_HUMAN, which is part of the train set, is an example of well performing SVD projection and neural mapping. The SVD predicted distributions of hits and nonhits are well separated and close to their original ranges. The NN mapping is not perfect but results in a distribution where the hits and non-hits have clearly identifiable modes. The kinase KIT_HUMAN, which is part of the validation set, performs almost as well. Note that SVD mapping is based on the *training* kinases, since only those are included in the set that defines the SVD basis. The imperfection of the SVD mapping for these kinases is simply due to truncation.

By contrast, the kinase vectors for the *test or validation set* are not included in the construction of the SVD basis, and we expect some loss of accuracy even with perfect SVD mapping (*N*_2_ = *N*_B_).

The three kinases shown that are part of the test set illustrate qualitatively different situations. For the kinase DYR1A (right column), a significant fraction of hits is mapped by the SVD truncation to above the support of the non-hit distribution. These hits are then mapped by the neural net quite faithfully, so that many hits are in the top of the predicted affinities. These hits can be easily recovered. By contrast, many of the hits of the kinase PGFRB and KIT are mapped already at the SVD step into the distribution of the non-hits. The NN mapping only adds some noise, and the bulk of the cloud of hits is embedded in that of the non-hits; yet the resulting distribution can still be used to extract a set of ligand groups that are enriched with hits.

As it should be obvious from the subsequent analysis, the qualitative outcomes of the SVD and NN mappings are not exclusive to one or the other of the train, validation, and test sets of kinases, albeit the train set kinases perform better, on average. A meaningful comparison should consider measures of performance and evaluate them for all kinases.

### Evaluation of results

Ideally, when examining the distribution of predicted affinities, the values for hits (known interacting ligand groups) for a kinase would form a distinct mode, clearly separated from that of the non-hits. The ranges and mean values of these distributions for each of the 455 documented kinases are depicted in Figure 5. Although the values corresponding to hits are generally higher, the ranges typically overlap, indicating some level of ambiguity in distinguishing hits from non-hits based solely on predicted affinity.

**Fig. 5:**
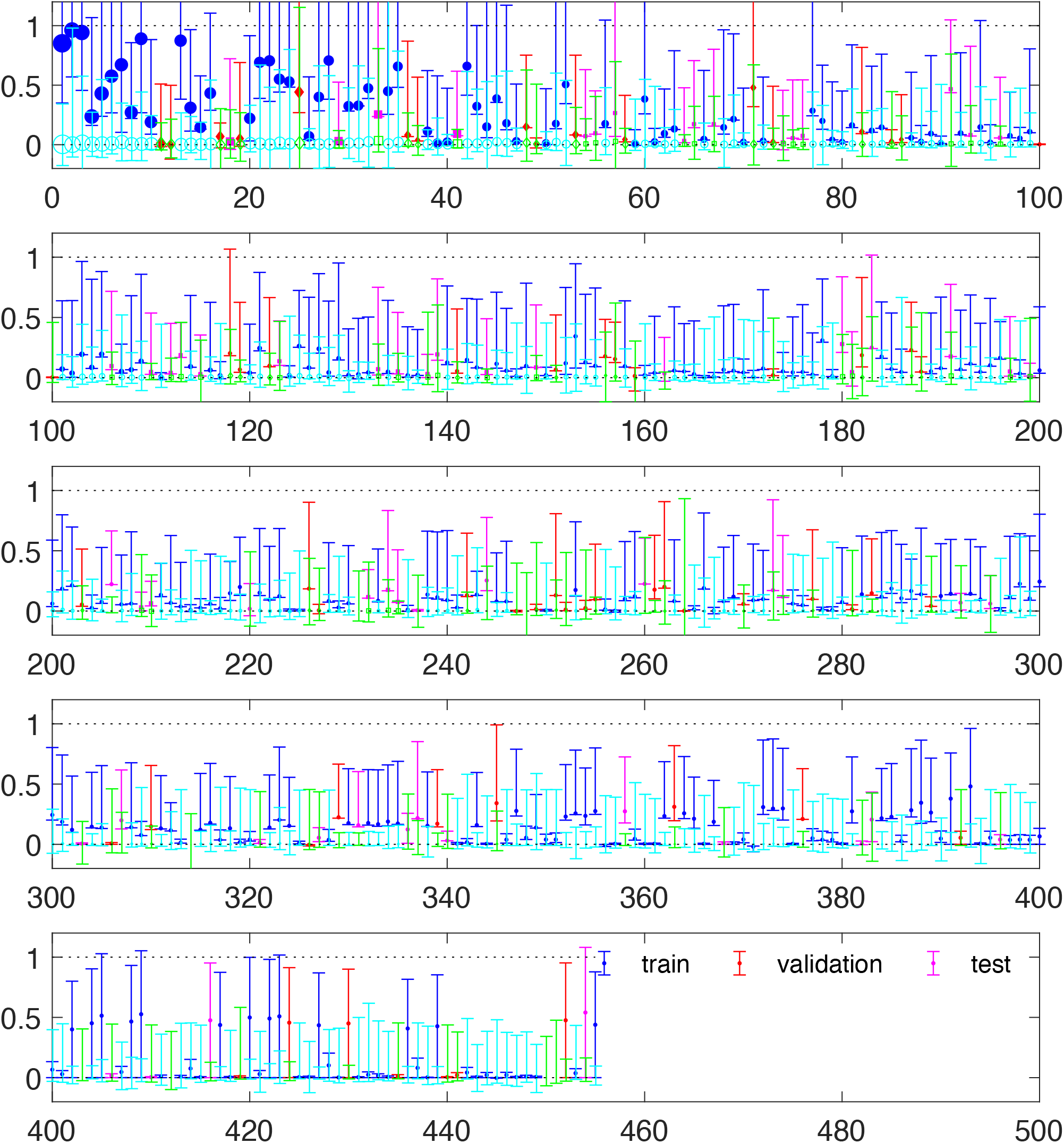
Ranges and means of the distribution of predicted scaled affinities for the hits and nonhits of each of the 455 kinases, from one model run. The ranges typically overlap but the hits are higher than the non-hits. Here, kinases are sorted by the number of known interactions (hits) for the respective kinase, also indicated by the sizes of the solid circle markers. Train, validation, and test groups are indicated by color. (This plot represents results from run #74839.)

The challenge of identifying hits using a threshold value for their predicted affinities represents a classic binary classification problem, where the task is to separate positives and negatives based on a single quantitative measure (DeMaris & Selman, 2013).

We computed Receiver Operating Characteristic (ROC) curves for individual kinases, as well as for combined sets of kinases. The area under the curve (AUC) from these ROC curves is a widely utilized measure of performance. The distribution of AUC values for individual kinases is illustrated in Figure 6.

**Fig. 6:**
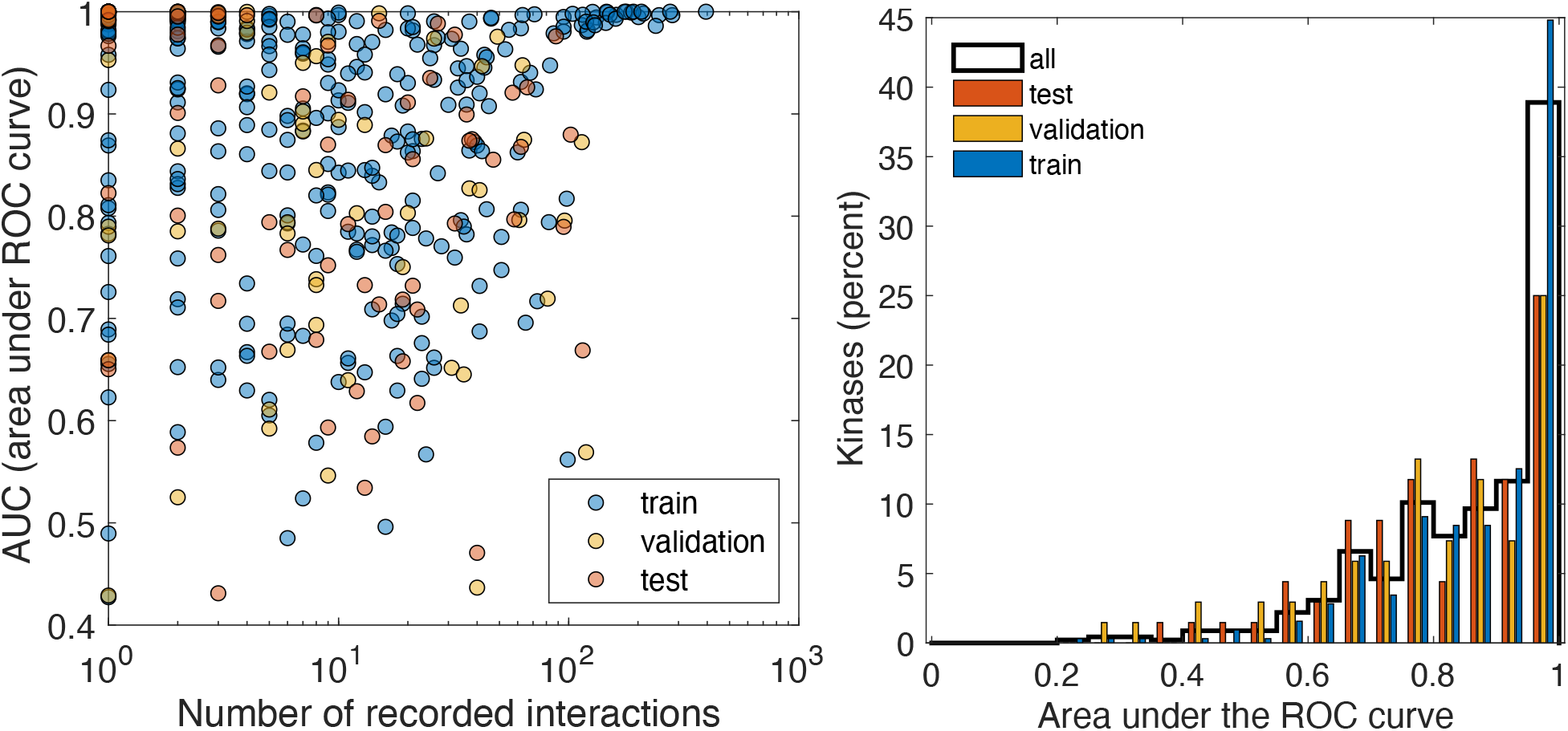
Area under the curve (AUC) estimated for individual kinases using the respective receiver operating characteristic (ROC) curves. The left panel shows a scatter plot of the values, with AUC on the y axis and the number of known hits for the respective kinases on the x axis. Train, validation, and test sets are indicated by color. A histogram of the AUC values of the three kinase groups is shown in the right panel. The test set performs worse than the train set (as expected), but the majority of kinases in the test set have AUC above 80-90%, indicating that the predicted affinity values are a good predictor. (This plot represents results from run #74839.)

The average AUC by kinase (for this model run) is 87.79% for the train set and 81.44% for the test set. However, if we weight the kinases by the number of hits, the average AUC works out to 93.34% for the train set and 81.90% for the test set. Computed globally (ROC curves for all hits from all kinases in the respective set) it works out 93.64% (train) and 82.76% (test).

These ROC-derived measures do not fully capture the challenges arising from the fact that the number of hits is typically very small compared to the total number of ligands. The number of hits per kinase can range from as few as 2-3 up to a maximum of about 350 (in the case of VGFR2), compared to a total of 4500 − 4800 total ligands accounted for in a model run with typical training / validation set of kinases (4731 for the run discussed throughout this section). The mean (27.82) and median (9) number of hits per kinase from the 4731 ligands represent 0.59% respectively 0.19% of the total.

The issue is illustrated in Fig.S4. The ROC is based on the true positive and false positive *rates*, numbers of positives and negatives that fall above a given cutoff as a fraction of the total number of positives respectively negatives. A comparison between the absolute counts of positives and negatives (right panel) reveals that false positives can overwhelm the true positives even if the former represent a small fraction of the total number of negatives.

We also assessed the precision 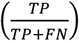 and recall 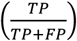 for the top 1% and 5% of ligands sorted by predicted affinity. We chose to use *relative* cutoffs – defined as a fraction of the total number of predicted affinities for a given kinase, rather than a specific cutoff value to be applied globally – because the range of the predicted affinities (for hits and non-hits) varies significantly between kinases. For example, in Fig.4, where kinases are sorted by the number of known hits, the range of predicted affinities is wider for kinases with many hits compared to those with only a handful. For the model run presented here^14^, using the top 5% of all ligands, the mean recall is 52.45% and the precision is 4.68% for the test set, compared to 59.07% and 9.66% respectively for the train set.

The relatively low precision is attributable to the fact that the proportion of true hits is significantly less than 5%. The uneven distribution of known hits among kinases (recall that 120 of the 455 kinases have 80% of the known hits) is likely to play a role here. We also computed the recall globally (sum of all recovered hits / sum of all hits); in this case we would expect the well-documented kinases (with many hits) to have a higher impact. This results in a clear difference in favor of the train set compared to the test set.

When the cutoffs are adjusted to retrieve 50% of the hits for each kinase, the average precision per kinase increases to 17.7% (33% for the training set). This indicates that while the model is reasonably effective at identifying a broad set of hits, the large number of potential non-interacting ligands challenges the model’s ability to pinpoint hits with high precision.

**Table 1.**
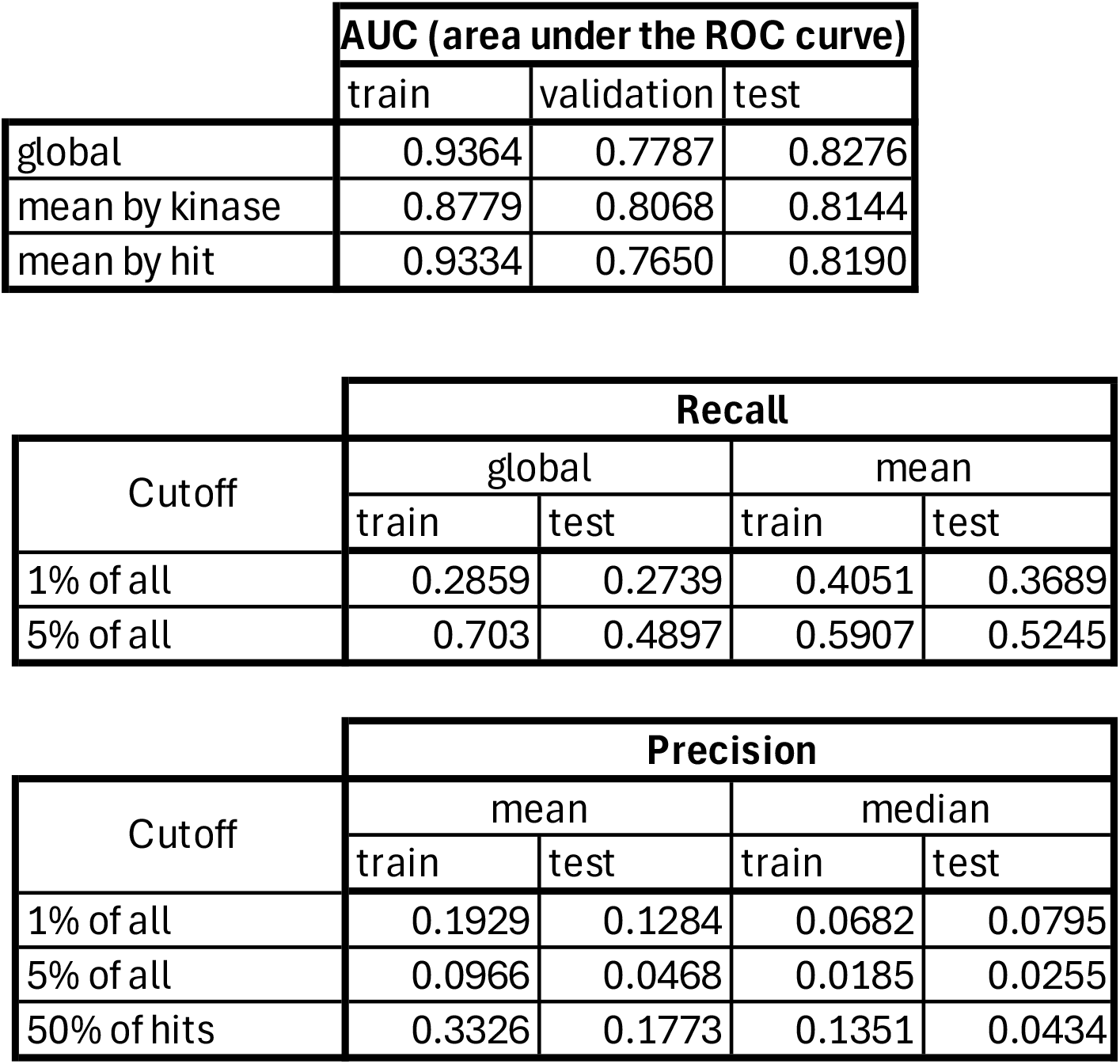
Values obtained for the AUC, Precision and Recall measures, for one model run (ID 74839, Set 2). Values labelled global were obtained by counting all the hits or non-hits of the respective kinases, above a given cutoff or in total. For the recall, this is equivalent to a weighted average of the by-kinase recall values. (Results from run #74839.)

### Linear Regression and Combined Models

#### Linear Regression Models

We also developed a linear regression-based model framework for the kinase structure / activity dataset. This model duplicates the major steps of the neural net approach as described previously, except for the prediction of coordinate vectors (PCA components of the molecular structure vectors as input, to PCA components of ligand affinity vectors), which is done by multiple linear regression instead of a neural net. The linear regression calculations are performed independently for each output PCA component, using the same (sub) set of input PCA components. We generated instances of the linear model using subsets of the first 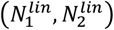 PCA components of the input respectively output coordinate vectors, mirroring the NN approach.

#### Neural Net versus Linear Regression Performance

A comparison of the performance measures resulting from the linear regression model with those from *comparable* neural-net models is shown in Figure 8 and S5. The typical NN performance is the highest for *N*_2_ *≈* 20 … 30, declines afterward, and increases slightly as the maximum output dimension (*N*_2_ = 289) is approached. The NN model consistently outperforms linear regression for the same input / output dimensions (when 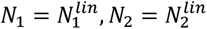) for output sizes of *N*_2_ ≤ 100.

The highest NN performance for this choice of train and test sets^15^ was AUC = 0.8276 for the train set, obtained with a NN model with *N*_1_ = 100, *N*_2_ = 20 and hidden layer sizes (1000,500,200). The mean performance for NN models with the same input and output sizes was AUC = 0.7975 (9 model runs). The linear regression model performance for <u>the same</u> input and output sizes 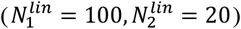 is respectively AUC = 0.499 and AUC = 0.5378 for 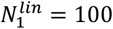 and 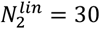.

However, the linear regression model performance increases consistently with 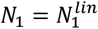 and 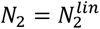 and overtakes the NN performance for *N*_2_ > 200. The highest overall performance of the linear model has been AUC = 0.8261 (mean: AUC = 0.8053) for 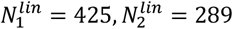, obtained when including the complete coordinate vectors (all PCA components) on both the input and output side. When restricting to 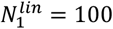, the top linear regression performance is AUC = 0.8155 obtained with 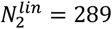.

In summary, we find that the MLP based neural net model outperforms the linear regression model when used with the same moderate input 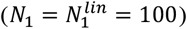 and low to moderate output dimensions 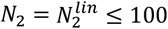. On the other hand, linear regression performs better than the NN for *N*_1_, *N*_2_ *≈* 200 or higher.

#### Combined NN / Linear regression

We constructed combined predictions from pairs of specific NN and linear regression models, by assigning a the higher of the two predicted scaled affinities for each kinase and ligand group. Any two model prediction sets can be combined in this manner; we constructed combined models from a given NN model by either adding a linear regression model with the same input and output sizes (“linear match”) or by adding a full-size linear regression model (“linear max” where maximal sets of PCA components are used).

**Table 2.**
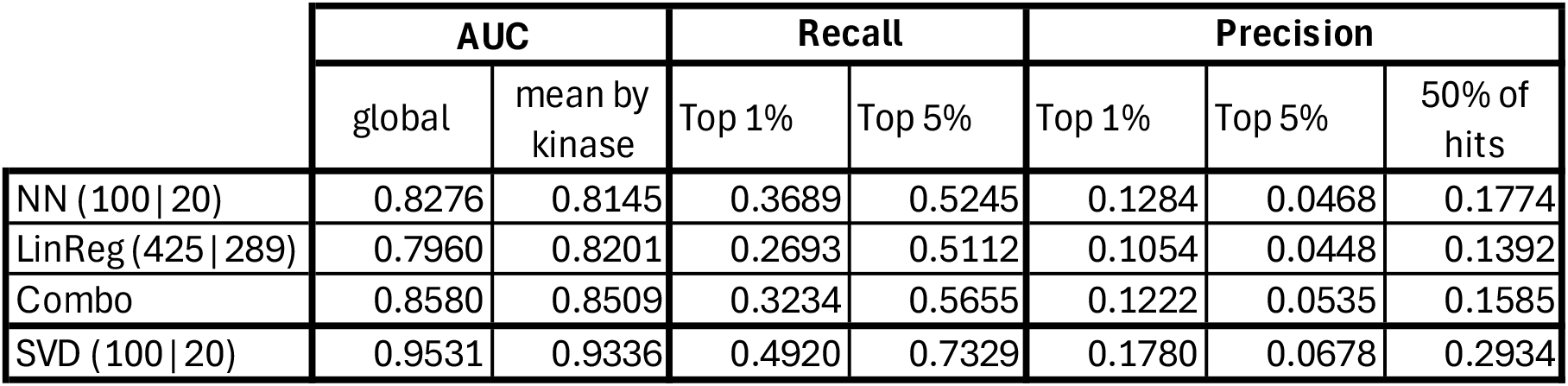
Comparison of performance measures obtained on the training set for the AUC, Precision and Recall measures, for one NN model run (ID 74839 as described previously); linear regression using all PCA components, and their combination.

The combined models typically provided the best performance, with top AUC scores above or approaching 0.87. For instance, combining the above described NN model output with that of a linear regression model using the full sets of PCA vectors, yields global AUC = 0.8509 which supersedes both the NN and the linear regression scores. This is typical in some parameter ranges but is not always the case; the combined scores sometimes fall between the original ones (but never below both of them, as far as we checked). Such instances are illustrated in Figure 7 and S5, where the performances of various model combinations are shown as a function of the output size.

**Fig. 7:**
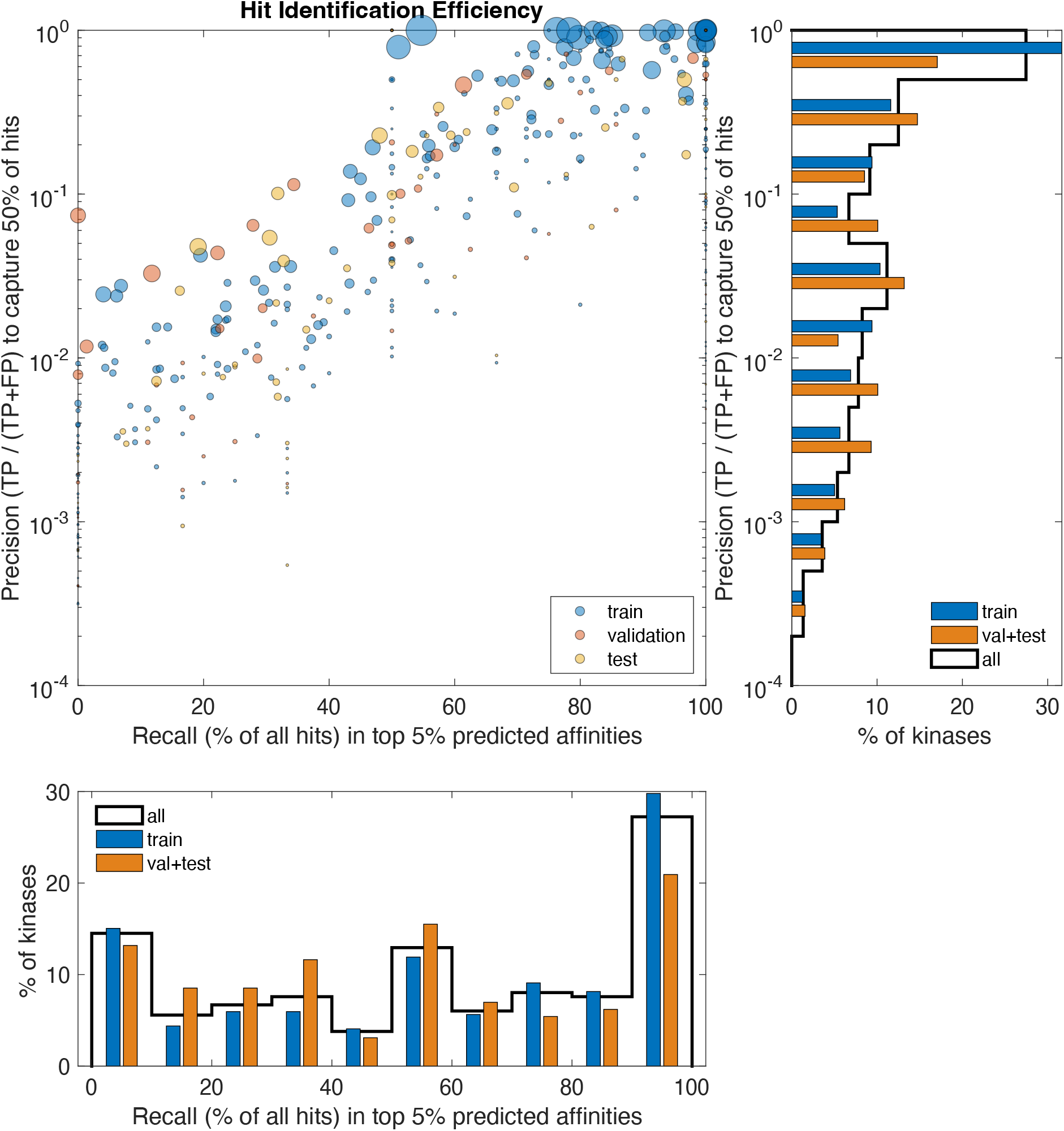
Precision and Recall measures obtained using the NN model for one model run (#74839). Precision is calculated for individual (by-kinase) cutoff values that retrieve 50% of the known hits*; Recall is calculated using individual cutoffs at the top 5% of all predicted affinity values* for the respective kinase. Train, validation, and test sets are indicated by color. Marker sizes indicate the number of known interactions (“hits”) for the respective kinase. (*Ligand groups that do not interact with the train set kinases are omitted from the hit count of test and validation kinases). Histograms on the sides are marginal distributions and represent the number of kinases with the respective performance measures. (This plot represents results from run #74839.)

**Fig. 8:**
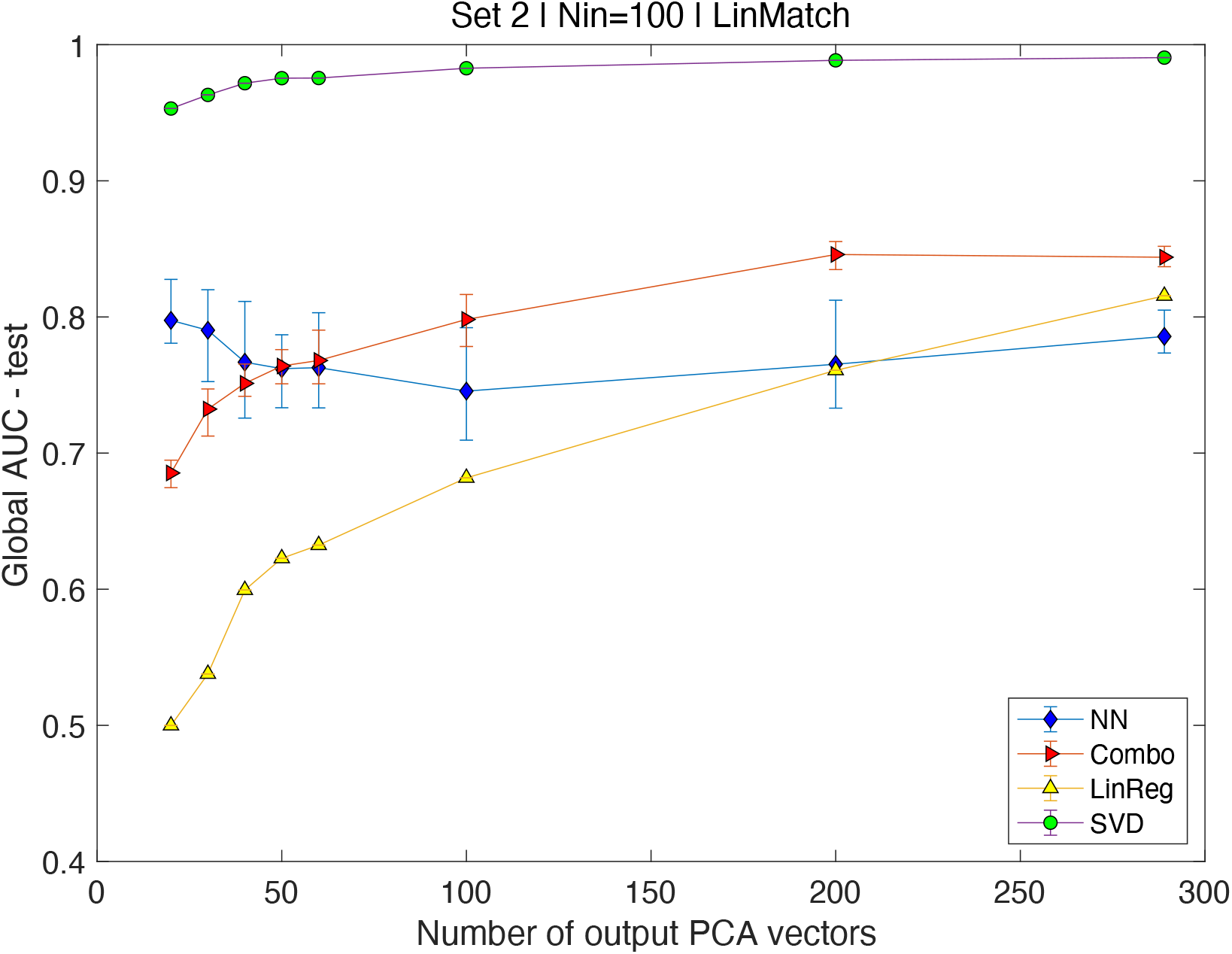
Summary of a performance measure (area under the global ROC curve) for multiple model runs, including NN, linear regression, and combined. All model runs have been performed with the same selection of training (319), validation, and test kinases (68 each); the PCA basis for the output vectors (ligand group affinities) is constructed using the training kinases only, and ligand groups that do not interact with the training kinases are excluded.

## Discussion

### Overview / context

Machine learning (ML), particularly within the realm of artificial intelligence (AI), has significantly impacted drug development, notably through AI-assisted approaches to protein folding and molecular dynamics studies (Abramson et al., 2024). However, the task of predicting binding affinities from molecular structures, based on interaction energies associated with the geometric arrangement of molecules, remains resource-intensive. The vast space of potential candidate molecules and binding configurations renders exhaustive (“brute-force”) calculations prohibitively expensive, even with today’s available resources. Therefore, there is substantial value in identifying likely ligand candidates for a specific molecular target using a data driven approach. Our approach leverages annotation-level data about targets and their known interactions, along with limited structural information about potential ligands, to identify ligand-target pairs using deep learning. The data we used was mined from Pharos / TCRD, which aggregates available research data on human proteins (Kelleher et al., 2023) (Oprea et al., 2024).

### Data source and organization

#### Kinase information (ligand-independent)

We utilized data on 455 kinases with recorded ligand binding information in TCRD to develop a deep learning model with the goal of identifying novel (yet unknown) ligands for all 635 kinases. This regression model predicts scaled binding affinities for specific targets concerning a set of ligands, based on standardized information about the targets’ molecular structure (molecular fingerprints) and gene ontology annotations. The regression model is designed to use this information, formalized in binary vectors, to predict binding affinities between targets and ligands.

#### Ligand binding information

We employed existing ligand binding information to train and test deep learning and linear regression models. Through a user interface (Pharos) query, we retrieved *≈* 106,518 kinase-ligand interactions involving *≈* 58,798 ligands and 455 kinases, counting all interaction types and some multiple sources for the same interaction. These involved *≈* 81,000 distinct kinase-ligand pairs. We characterized the interactions between any kinase-ligand pair with the highest recorded affinity regardless of interaction type. Arranging into ligand interaction vectors by kinase results in a sparse 455 × 58,798 matrix with *≈* 81,000 nonzero entries.

### Modeling strategy

#### Ligand clusters

To reduce the dimensionality of the output vector we clustered the ligands into *≈* 5275 groups using a structural similarity measure. We constructed a matrix of kinase-ligand group affinities by assigning the highest affinity between a given kinase and any member of the corresponding ligand group. This 455 × 5275 matrix, with *≈* 50,000 entries, was the primary data used to develop the deep learning (and linear) regression models.

#### Regression in coordinate space

Our regression model outline is illustrated in Figure 1. The 2770, respectively 5275-dimensional input and output vectors are projected onto SVD bases of their span. Neural-net and linear regression models are trained on the corresponding pairs of *coordinate vectors* (that consist of the coefficients used to reconstruct the original vectors). This provides a way to reduce the dimensionality of the input and output sizes of the regression models to (at most) the dimension of the respective spans^16^, 425 and 289, respectively. Further dimensional reduction follows from using truncated coordinate vectors, using only the first *N*_1_ ≤ 425 respectively *N*2 ≤ 289 components of the coordinate vectors. This *truncated representation* comes at the price of less than faithful reconstruction of the original ligand affinity vectors, even if the regression model outputs were perfect.

We developed and trained multi-layer perceptron (MLP) models using combinations of (*N*_1_, *N*_2_) dimensional input-output vectors pairs corresponding to a subset of the 455 documented kinases. We found that the NN model performance was at its best for moderate size networks with 100 − 200 input dimensions, 20 − 40 output dimensions and three hidden layers with sizes ranging from 500 to 2000. While the model performance improved as the output size approached the maximum, the reduced performance is very likely the result of challenges / logistical limitations in training the larger models.

#### Parallel Neural-Net and Linear Regression models

In parallel with the neural-net based models, we developed a type of linear regression model trained to predict the same pairs of coordinate vectors. Due to its simplicity, this model can be trained more easily, and its performance increases with the number of included coordinates (principal vectors). For the range where the NN model works well (≤ 50 input dimensions), the NN outperforms the linear one. As the number of dimensions increases, the linear model performs better. However, combined models where the predicted affinities are taken to be the higher one of the two predictions, provide the best performance.

### Classification Performance

The primary model output consists of (predicted) scaled ligand group affinities which can be used to identify likely ligand groups that will interact with a given kinase (target). Identification of interacting kinase-ligand group pairs is a *binary classification* problem, aiming to identify hits (reacting ligand groups for a given kinase) by choosing a cutoff value for the predicted affinity. We computed several performance metrics for multiple instances of the regression models.

The area under the ROC curve (AUC) is a measure of the overall performance when using a continuous quantity of interest to identify positives and negatives^17^. A value of AUC = 1 represents perfect separation between negatives and positives while AUC = 0.5 is equivalent to random selection. We also computed Precision (true positives / all predicted positives) and Recall (true positives / all actual positives) measures corresponding to specific cutoff values. We relied on cutoff values defined in terms of a fraction of the total, at the top 1% and 5% of all predicted affinities for a given kinase or of all kinase-ligand group pairs for multiple kinases. These measures would be applicable to kinases for which ligand binding information is unavailable or incomplete.

We illustrate the results from a model run (# 74839) with 100 respectively 20 input and output dimensions and hidden sizes 1000|500|200. The global AUC (all test kinases) was 82.86% and the average by kinase was 81.44%. With a cutoff at the top 5% of all active^18^ ligands, recall (mean by kinase) was 52.45% and precision was 4.68%. A more selective cutoff of 1% yields a recall of 36.89% and precision 12.84%.

The low precision values reflect the fact that the cutoff percentages of 5% and 1% exceed the total number of hits of many of the kinases. The total number of active ligands is 4500 − 4800 (depending on the choice of the train set; it is 4731 for the model run discussed here), with ∼250 respectively ∼50 ligand groups in the top fractions. By contrast, the number of hits per kinase is 27.8 on average, but many (about half) of the kinases have 10 or fewer hits. On the other hand, some kinases do have many hits, so the 5% and 1% cutoffs will limit the recall performance. The precision corresponding to a cutoff at 50% of the known hits is a measure that gives a more balanced picture of the ability to retrieve hits based on model predicted affinities^19^. For the model run discussed here, we have 17.7% for the test set (compared to 33.2% for the train set).

### NN Performance Plateau

We explored a range of multi-layer NN models by considering more hidden layers and larger input, output and hidden layer sizes, as well as multiple selections of the train, validation, and test kinases. Keeping the number of hidden layers fixed, increasing the size of one or more of the layer sizes eventually led to diminishing returns in terms of performance, in addition to the obvious increase in computational cost. Figures 8 and S5 show the dependence of one measure (global AUC) on the output size (number of PCA components used), while keeping the number of input vectors fixed.

Typically, the performance of the NN models peaks at *N*_2_ = 20 … 30, then diminishes for intermediate values 50 … 200 and finally rebounds approaching the top values as *N*_2_ approaches the full rank. The output dimension, equal to the number of PCA components used in truncated representation of the ligand group affinity vectors, is the parameter with the most important impact. It is counter-intuitive but not surprising that the performance of a model with *N*_2_ = 30 cannot be matched by training a model with *N*_2_ = 60, even if we increase the sized of the hidden layers. It is likely that this is a convergence issue, in that the optimization algorithm is less effective in the larger search space; if that is the case, this can be likely resolved by more careful algorithm selection.

We investigated the loss of predictive power resulting from this truncated representation and found that the impact varies greatly from one kinase to another. Presumably this goes back to the way the SVD basis is constructed, with some kinases having more of their weight carried by the first few vectors, while other kinases (for instance, those interacting with rare ligands) being carried by basis vectors that have a small singular value.

### Linear Regression

We devised a linear regression model for the same sets of coordinate vectors as those used for the NN models. While the NN model is trained for all output dimensions simultaneously, the linear regression model consists of *N*_2_ separate linear regressions that are trained with all *N*_1_ input dimensions. Because of this, and the simplicity of linear regression, the training process is successful for larger input and output dimensions as well. Thus, as illustrated in Figs.7 and S5, the performance of the linear regression increases as the sizes increase, reaching top performance when *N*_1_ and *N*_2_ are maximal.

While the NN models for small to moderate dimensions (*N*_2_ ≤ 50 … 100) outperform linear regression models for the same input and output sizes, full-size linear regressions match the performance of the best NN models, reaching, AUC *≈* 0.826 (on average AUC = 0.805).

### Combined LR and NN models typically have the highest performance

We combine NN and linear regression models by building predicted affinity vectors whose entries are the higher of the corresponding entries in either model. These combined models typically outperform *both* input models and result in the best performance measures, reaching AUC = 0.879.

### Conclusions and Outlook

Machine learning driven approaches, particularly Deep Learning, can identify complex relationships, often matching or surpassing mechanistic models where such insight exists and providing predictive power even where mechanistic insight is lacking. In the context of target-ligand binding, there is obviously a causal link between the target and ligand structure (protein sequence and resulting 3d structure) and their binding capability. However, the causation traverses many layers, and first-principles approaches face challenges due to associated complexity. In protein-ligand docking scenarios, one only needs to find the minimum of the potential energy, but the conceptual simplicity is deceptive, since the energy function defines a complicated landscape in a very high-dimensional space.

The performance values we obtained are encouraging. For the DL models, the recall of 52% for a cutoff at the top 5% of predicted affinities (for the test set) signifies a more than ten-fold enrichment; the precision is 4.68%. With a 1% selection, the recall is 36.9% and the precision increases to 12.8%. Performance is further increased by combining with linear regression. While linear regression outperforms the NN for larger input and especially output dimensions, there is strong indication that the two model types provide complementary information; for moderately large sizes, the combined model outperforms both component models. The better performance of LR for high dimensions likely indicates the importance of including all SVD components, but the higher performance of the DL model for low dimensions indicates that the LR model cannot extract all the information carried by the respective first few SVD vectors. Therefore, a DL model along the lines we presented here, but with a more robust training and optimization strategy can potentially combine the benefits of both dimensionality and the ability to reproduce nonlinear functional dependence.

The performance level achieved here is likely to be significant for drug discovery applications as it suggests that selecting a subset of 5% of the ligand groups from the 5,275 clusters we identified will contain close to 50% of the “hit” groups, thus resulting in a 10-fold enriched set of potential ligands. This baseline can readily be improved by combining the DL model with a linear regression model trained on the full set of PCA components.

The performance of the LR model is intriguing, as is the fact that it seems we can predict or gain insight into the likelihood that a kinase will interact with a type of ligand, based only on human-generated, *higher-level* annotation-type information on its structure and gene ontology. This suggest that the linear structure of the ligand vectors is quite closely disentangled by or mapped onto the SVD basis, and that corresponding structures exist on the input side, among the molecular structure and gene ontology features encoded in the input vectors.

Further improvements are likely to be achievable by alleviating the dimensionality aspects that likely plague the NN training. The training algorithms we used deliver optimal results for moderately sized networks, with diminishing predictive power for larger network sizes. This is likely an aspect that can be enhanced with more advanced network configurations and dedicated optimization strategies. Nonetheless, such limitations may persist due to the finite nature of computing resources. One dimension reduction strategy is the projection of large-dimensional vectors onto a basis (done via standard PCA here) and using the resulting coordinate vectors in the ML process. This reduces the dimensionality from several thousand components to the rank of the input and output matrices (425 and 289, respectively). Due to network size limitations (reflecting limited resources), we truncate the coordinate vectors to a smaller number of components. While the basis vectors from an SVD decomposition of the original vectors provide an effective approximation, the loss of detail due to truncation is evident.

We identified the effect of truncation on the predicted affinities, and at the network sizes we are currently using, a significant portion of the error (in reproducing known binding affinities, even for training vectors) is attributable to this factor. Therefore, the ability to use larger networks, better optimization algorithms, and a more careful choice of basis vectors is likely to improve the performance of our approach substantially.

In conclusion, this study demonstrates the potential of machine learning, particularly deep learning models, to predict ligand binding affinities and identify potential binding partners for kinases. By leveraging structural and gene ontology annotations along with advanced algorithms like multilayer perceptrons, we have shown that it is possible to predict interactions with a significant level of accuracy, even in the context of high-dimensional and sparse datasets. The use of ligand clusters and dimensionality reduction techniques such as PCA has enabled us to manage the computational challenges posed by the vast space of potential interactions. Our results underscore the value of machine learning in drug discovery, providing a pathway to efficiently sift through large datasets and highlight promising targets for further experimental validation. Moving forward, enhancing model performance through optimization and addressing the limitations identified will be crucial in refining our approach and achieving even greater predictive power in the identification of kinase-ligand interactions.

## Footnotes

1 These 455 kinases have at least one ligand interaction documented in TCRD and are categorized as Tclin or Tchem. The remaining 180 kinases, categorized as Tbio or Tdark, have no recorded ligand interactions.

2 For example, if *EC*50 = 0.5 *μM* = 5 × 10^−7^*M*, the corresponding affinity is *a* = − *log*_10_(*EC*50) *≈* 6.3. A higher affinity corresponds to a lower effective concentration (less of the ligand required for 50% maximum effect).

3 The PCA decomposition for ligand affinity vectors is based on the training set only, since the affinities are not known for the kinases for which we aim to make predictions.

4 The same vectors were used to build linear regression models, discussed further below.

5 That is, using all *N*_1_ input components *σ*_*k*,𝓁_ ; 𝓁 ∈ {1, … *N*_1_} as predictors for each sample output ξ_*k*_,_*j*_.

6 Here TP is short for “true positives”, TN for “true negatives” (hits, respectively non-hits predicted correctly), FN for “false negatives” (hits predicted to be non-hits) and FP for “false positives” (non-hits predicted to be hits).

7 At the lower end of the cutoff range *t* = *t*_*min*_ the curve goes through the point (1,1) since there are no predicted negatives: *FN* = *TN* = 0 → *TPR* = *FPR* = 1; at the upper end of the range it reaches (0,0) since there are no predicted positives *TP* = *FP* = 0 → *TPR* = *FPR* = 0. When the classifier works perfectly, i.e. there is a cutoff value *t*^∗^ where there are no false predictions, *TP*(*t*^∗^) = 1, *FP*(*t*^∗^) = 0, the ROC curve follows the upper edges of the unit square (0,0) → (0,1) → (1,1).

8 See for instance Fig. 5. Kinases with more hits typically exhibit a higher range of predicted affinity values for both hits and non-hits. This is likely an artifact of the normalization of SVD components.

9 Model runs are assigned an integer identifier chosen randomly. Run #74839 has 100 input and 20 output dimensions and three hidden layers with sizes 1000,500,200 respectively. The MLP model is trained on a subset of 319 of the 455 kinases with ligand binding information; the remaining kinases are split equally into validation and test sets (68 kinases in each).

10 The PCA basis for ligand (group) affinity vectors is constructed from vectors corresponding to the training set only.

11 Run ID #74839 with 20 output, respectively 100 input SVD components and hidden layer sizes {1000,500,200}

12 Only affinity values that are deemed significant are recorded as such in Pharos.

13 We use PCA, which first offsets the vectors by the mean vector, then performs SVD on the shifted input vectors. The basis in this case consists of the ortho-normal SVD principal components, in the order of decreasing principal value. The discussion here applies to other choices of offset and base as well.

14 Run ID #74839 with 20 output, respectively 100 input PCA components and hidden layer sizes {1000,500,200}

15 One of the 5 train / validation / test configurations, labelled “Set 2”. The best NN performance in terms of global AUC corresponds to run ID 74839.

16 The PCA basis on the output (ligand-group affinity) side is constructed from the *training set only*, since the output vectors are not available for the kinases for which the model makes predictions. The rank of the (319 ligand group affinity vectors in the) training set is 289.

17 In our case, the quantity of interest is the predicted affinity associated with a specific kinase and a ligand group, and the possibility of correctly predicting whether the pair does in fact interact (as defined by the existence of a non-zero known affinity).

18 Excluding ligand groups that do not interact with the training set.

19 Obviously, this measure can only be applied to kinases with known ligands.

