## supplementary figures for "Deep Learning enabled discovery of kinase drug targets in Pharos"

**Fig.S1:** Distribution of the number of documented ligand interactions ("hits") by kinase. Left panels are by ligand group, right panels by kinase. The bars show frequency (number of instances) – top panels in terms of kinases, bottom panels weighted by the number of known interactions of the indicated type. Of the 455 kinases with documented interactions, a significant fraction has very few interactions (133 have between 1 and 5); looking at the distribution hits, the vast majority of known interactions involve kinases with a higher number of interactions.

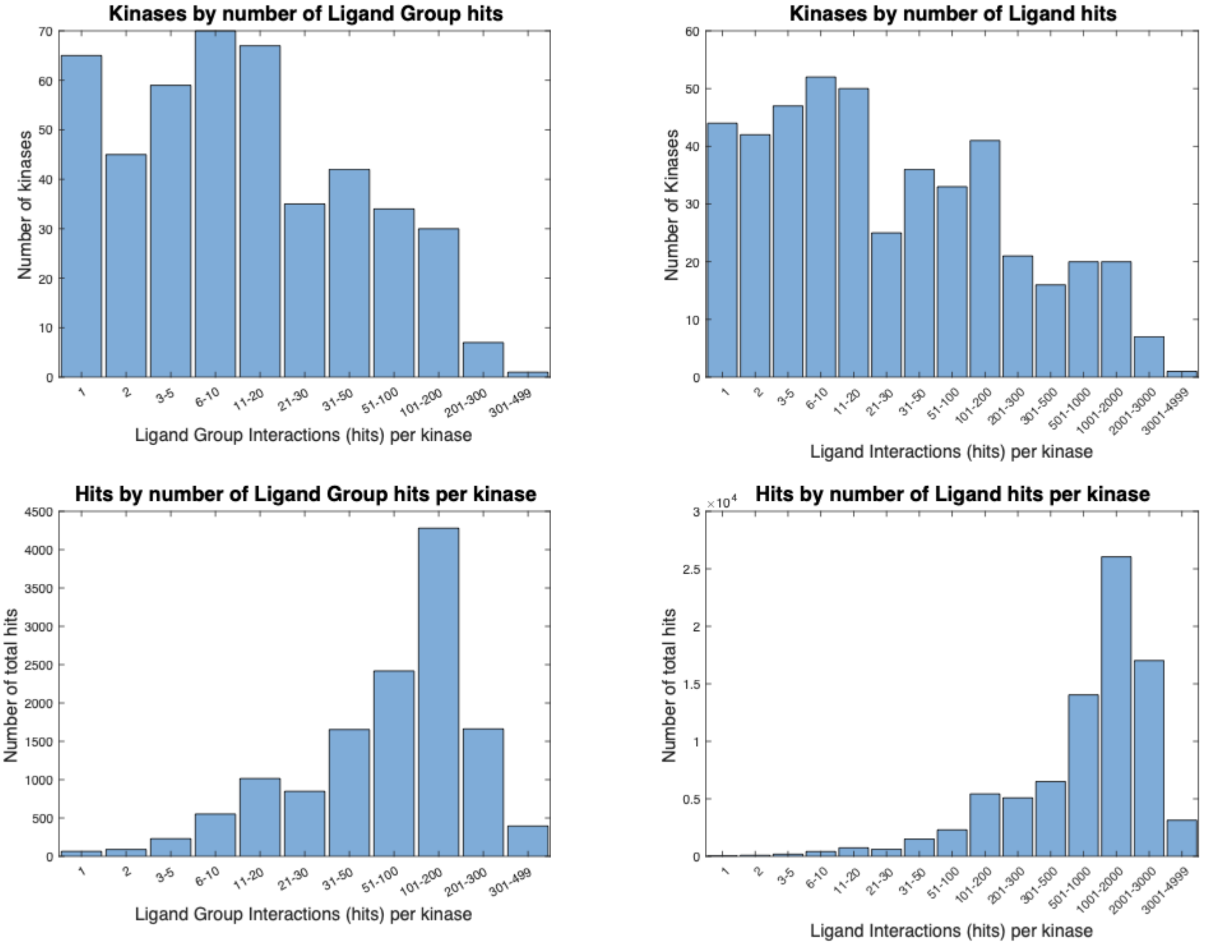


### Additional info

There are 106719 kinase-ligand activity entries in the original query file (LigandQuery.xlsx).

We keep only those entries with ligand SMILES (needed for ligand identification and clustering) as well as ligand activity type and values. This results in 106502 entries.

There are 58798 distinct ligand SMILES entries (in the remaining cleaned data) and 80878 distinct ligand – kinase pairs. These are the ligands and interactions included in the subsequent analysis.

**Fig.S2:** Distribution of the number of hits (known ligand interactions) by kinase, in terms of individual ligands (bottom) and ligand cluster or groups (top). Kinases are sorted by the number of known interactions in both panels. A minority of the kinases (64, respectively 120 of the total of 455) accumulate 80% of all known interactions. This is almost certainly a reflection of the research focus on these well-studied kinases.

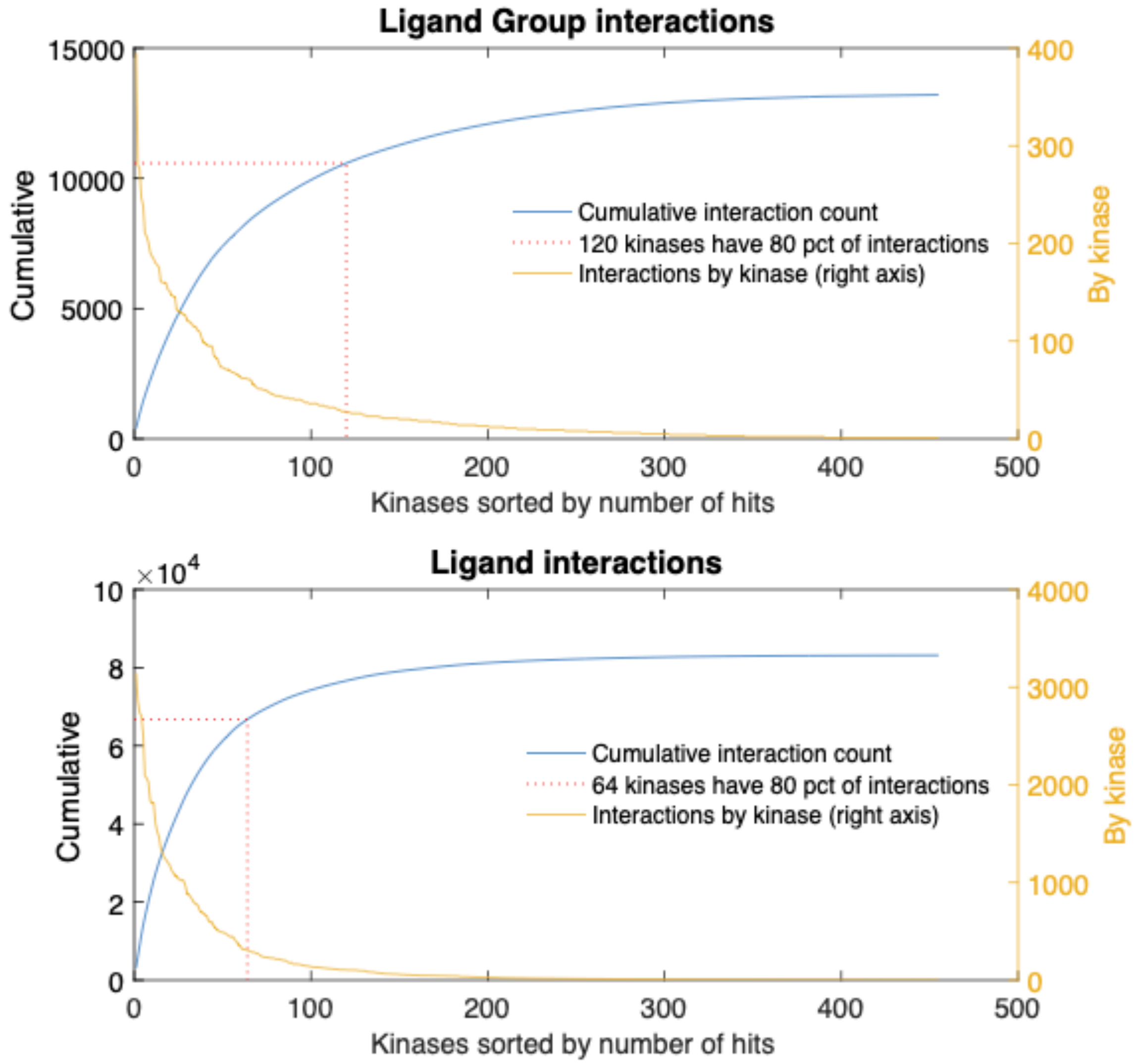

**Fig.S3:** The distribution of predicted affinities is a result of two mappings, one due to the truncated SVD basis and another due to the neural net. The bottom row of frames compares the neural net prediction against the SVD reconstruction obtained with the same number of PCA vectors. This comparison reflects the extent to which the neural net is able to reproduce the coordinate vectors (a perfect NN would result in a diagonal). The top row of frames shows the actual scaled affinities vs. the same y-axis. For some of the of the kinases the separation between hits and non-hits is lost at this level and could not be reobtained even with a perfect NN mapping.

The SVD basis covers the span of the training set of kinase affinity vectors but not that of the validation and test sets, thus the truncated SVD representation is on average more "lossy" for the test and validation sets. However, a wide range of overlap / separation levels are seen for all kinase groups. The loss due to projection decreases as the number of included components increases. (This plot represents results from run #74839.)

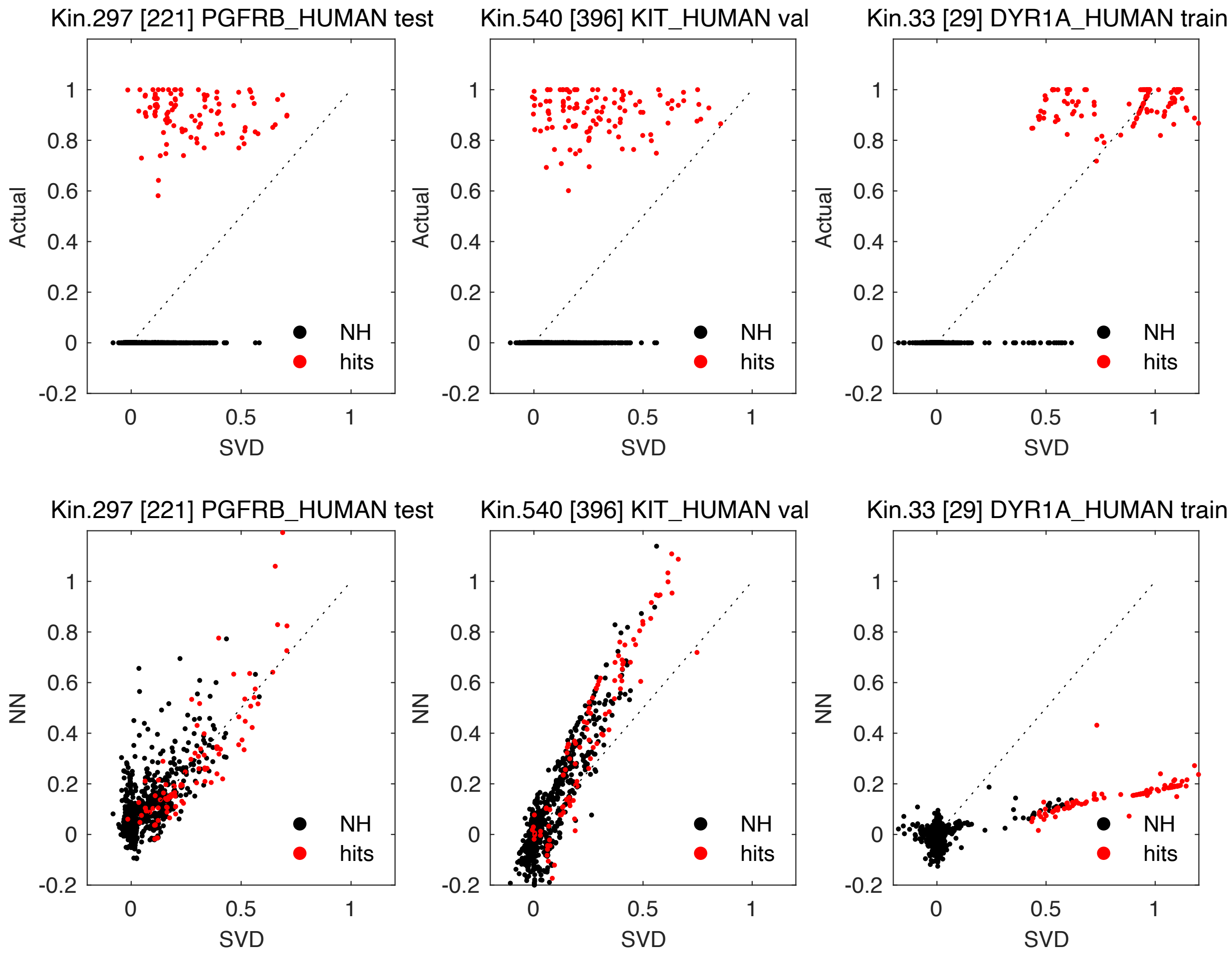

**Fig.S4:** Receiver Operating Characteristic (ROC) curves for the individual kinases (**right panel**). The area under the ROC curve is used as a measure of performance for binary classification; a value of 0.5 corresponds to random selection, and a value of 1 indicates perfect separation (non-overlapping distribution) between negatives and positives. (This plot represents results from run #74839.)

The ROC curve is a parametric curve defined by the relative numbers of positives and negatives that fall above a given cutoff compared to the total number of positives respectively negatives.

A comparison between the absolute counts of positives and negatives (**right panel inset**) reveals that the number of false positives can overwhelm the true positives, even if it represents a small fraction of the total. This is to be expected since positives are less than 1% (typically 0.1%) of the total. The inset plot shows the absolute number of true positives, false positives, and their difference (called the Youden statistic) plotted versus the cutoff values. While the TP counts hit a maximum of $\approx 12{,}000$, they only represent 0.6% of the total, since number of non-hits (counting ligands + kinase pairs) is $\approx 1.95 \times 10^6$.

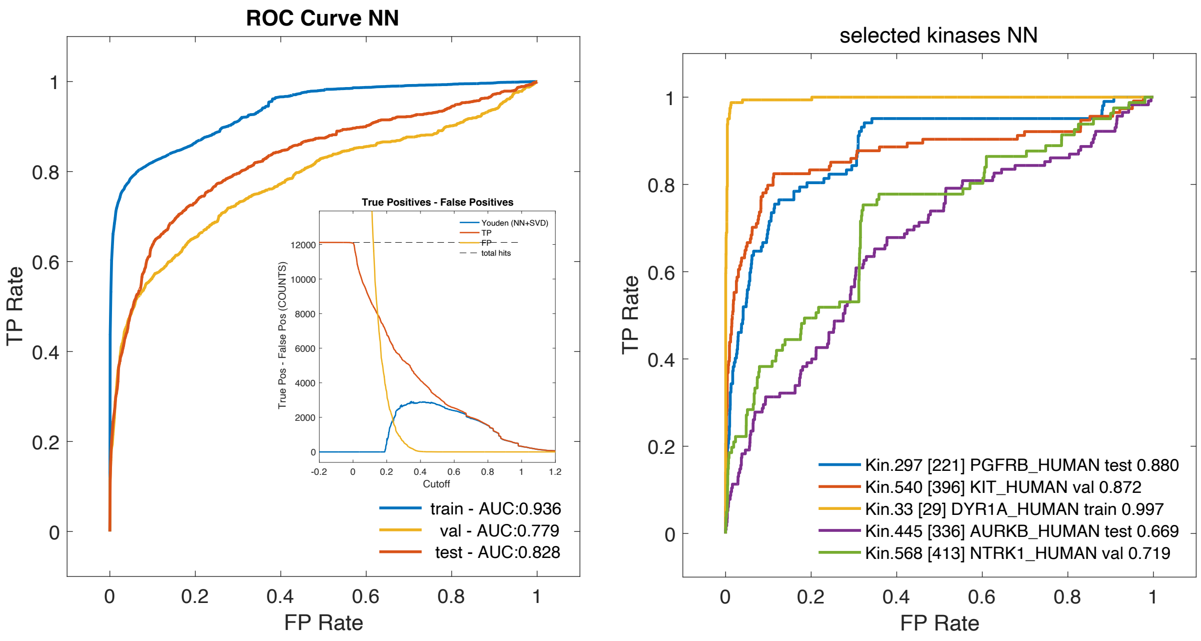

**Fig.S5:** Additional plots of the global AUC for groups of model runs. *Clockwise from the bottom left:* (a) Neural net (NN), linear regression (LR), and combined NN-LR models using $N_1 = N_1^{lin} = 200$ and $N_2 = N_2^{lin}$ as indicated on the *x*-axis – compare to Fig.7; (b) same as (a) with $N_1 = N_1^{lin} = 200$ for the NN and LR models and $N_1 = 200$ for the combined one, but the combination uses LR with maximum sizes; (c) same NN and LR models as in Fig.7 ($N_1 = N_1^{lin} = 100$), but the combination model uses a linear regression model with maximum sizes ($N_1^{lin} = 425, N_2^{lin} = 298$); (d) plot showing all NN, LR, and combination model groups to help appreciate the variations between the different subsets.

All panels shows the mean and range of the global AUC obtained from: (1) separate NN (blue diamonds) and (2) LR (yellow vertical triangles) models with the indicated output size ($N_2$ respectively $N_2^{lin}$ number of PCA components indicated on the *x*-axis); (3) combination models (red triangles) with the same input and output sizes as the NN models shown, but using LR models as indicated (when the NN and LR input sizes differ, the horizontal axis indicates the input size of the NN). Finally, (4) the performance obtained by PCA truncation only (corresponding to a "perfect" prediction of the $N_2$ components; green circles).

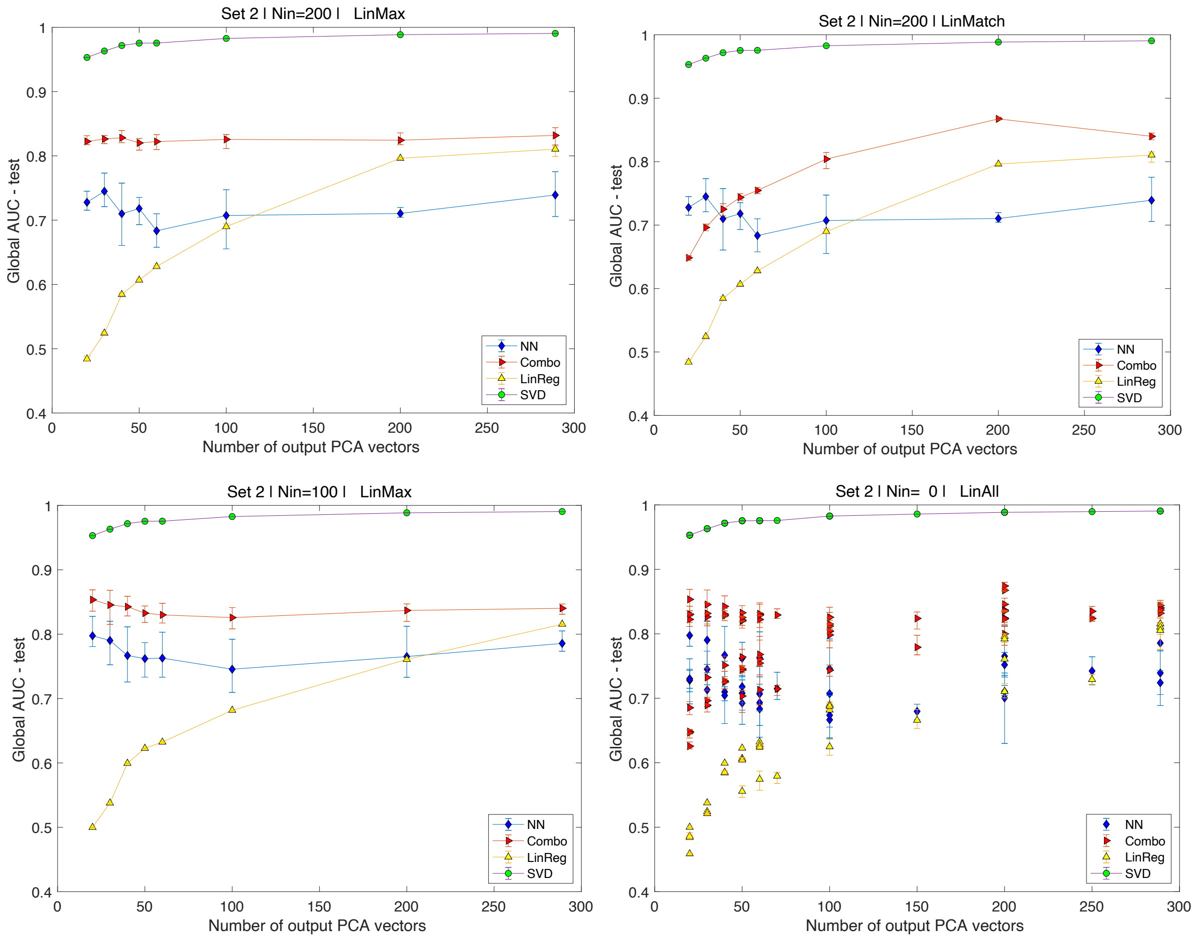
