## Supplementary material for "Deep Learning enabled discovery of kinase drug targets in Pharos": Figures and tables

**Fig.1**: Schematic of the multilayer perceptron-based model outlining the mapping $\vec{y}_{kin} = F(\vec{x}_{kin})$ from the 2770-dimensional input (structural + gene ontology annotation vectors) to the 5275-dimensional output (ligand group affinity) vectors. Both sets are projected onto a basis of their span* (obtained through PCA). The resulting coordinate vectors are truncated and only the first $N_1$, respectively $N_2$ components are used in the NN model. *The test set affinity ligand group affinity vectors are not included when constructing the projection (PCA) basis; ligands that only interact with the test set (with identically zero entries on the train and validation vectors) are excluded; the clipped test-set vectors are otherwise processed identically to the others.

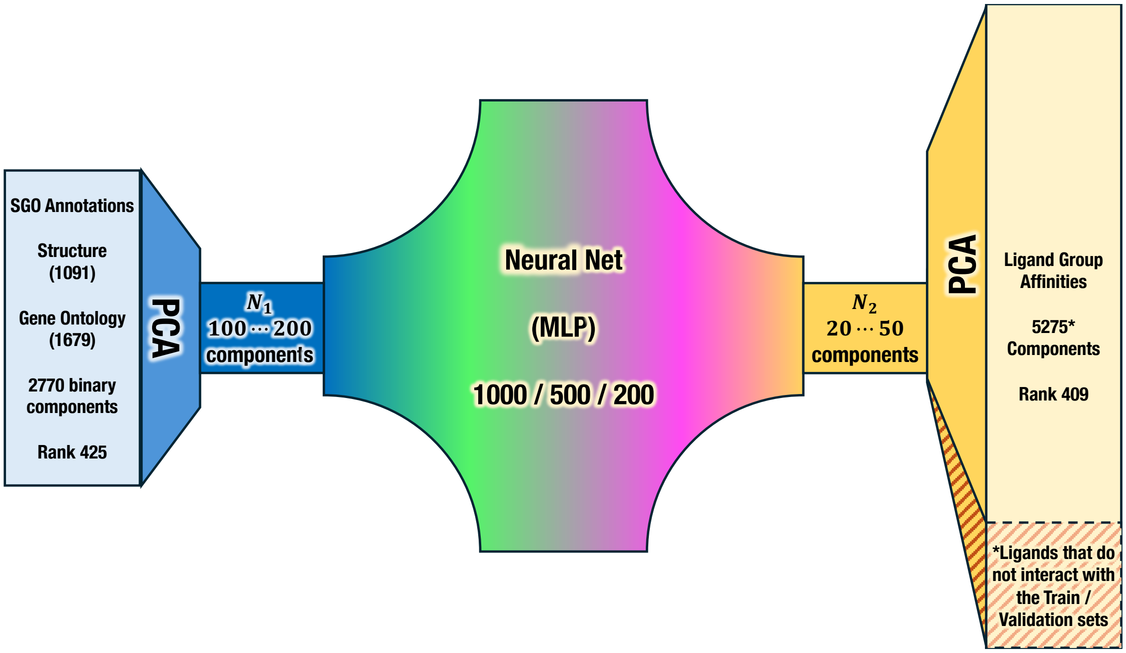

**Fig.2:** Distribution of the 80878 recorded (maximum) affinities of kinases to individual ligands. The shape of the distribution shows a significant drop below 7.3 ($\approx 30nM$), consistent with the policy followed in compiling Pharos / TCRD data. It is likely that many low affinity values resulting from laboratory measurements were not published.

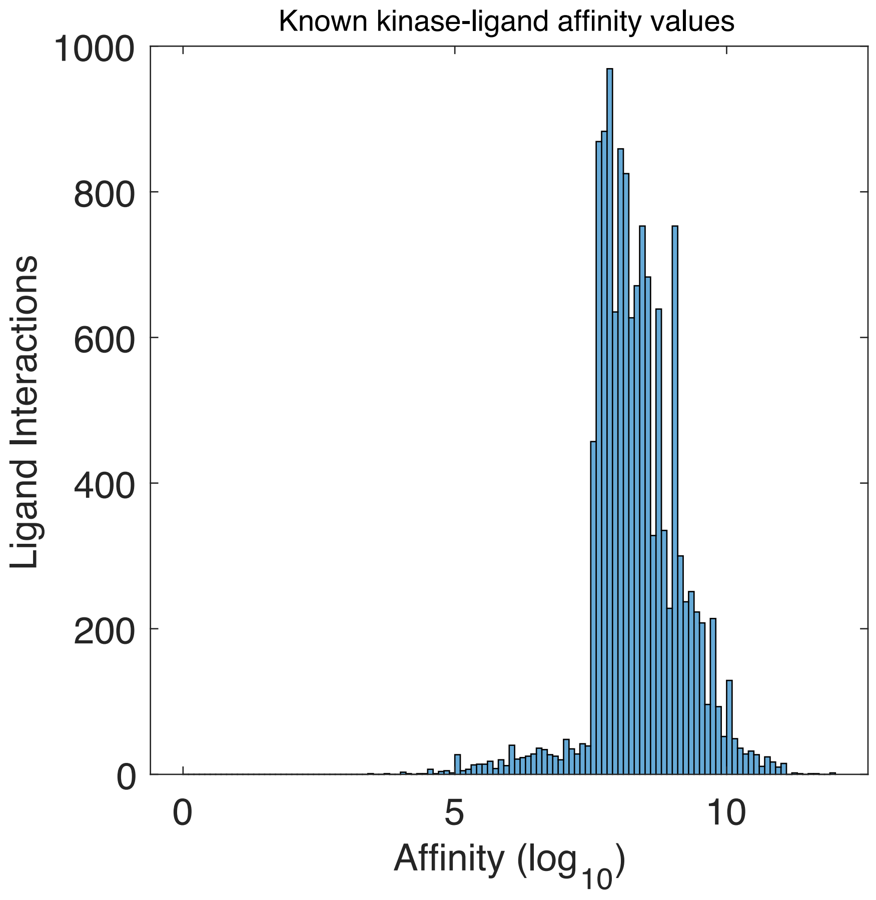

**Fig.3:** True and model predicted ligand group affinities for three kinases. The true affinities (left y-scale) used as training input are scaled to the [0,1] interval so that ligands with known interactions have nonzero affinities approaching 1; affinity values for ligands with no recorded interaction are exactly 0. The neural-net predicted affinities (y-scale on right-hand side) approximate the true values. For a given kinase, the predicted affinities induce a ranking of the ligand groups used to infer likely interactions. This is the basis of the performance measures discussed in the paper. Panels show different kinases; their role in the model training process is as indicated. This plot represents results from run[1] #74839.

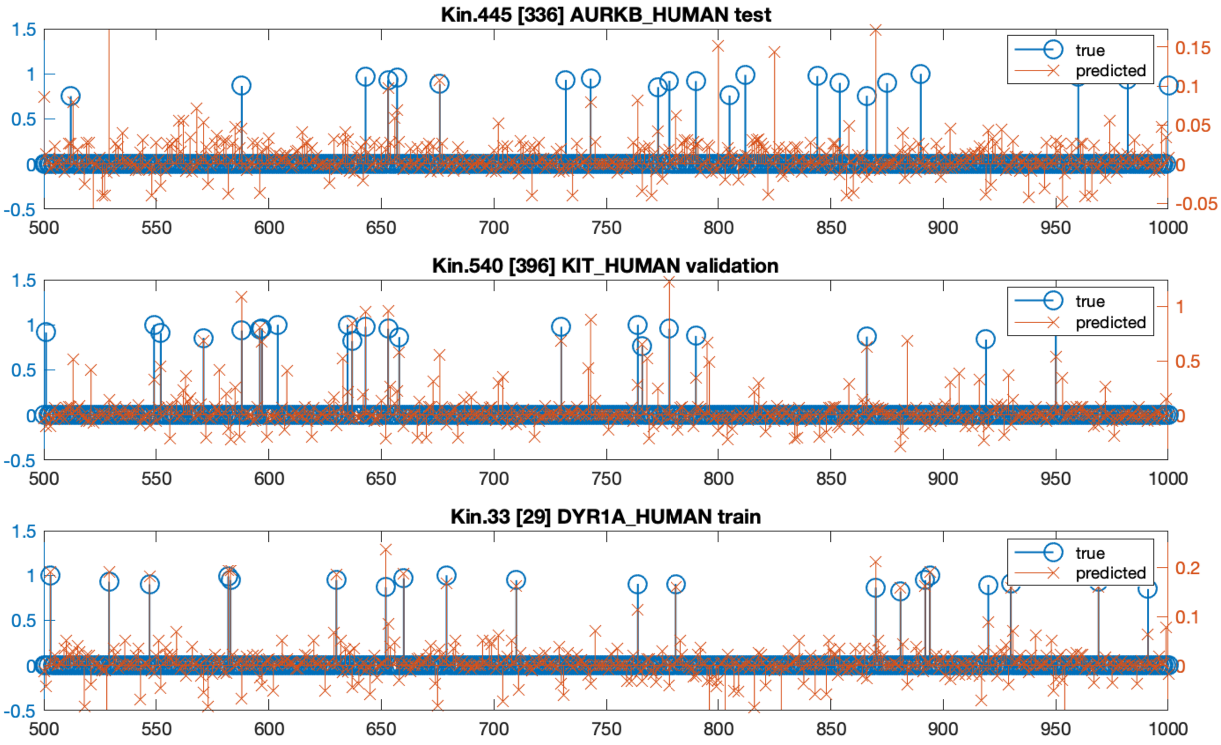


[1] Model runs are assigned an integer identifier chosen randomly. Run #74839 has 100 input and 20 output dimensions and three hidden layers with sizes 1000, 500, 200 respectively. The MLP model is trained on a subset of 319 of the 455 kinases with ligand binding information; the remaining kinases are split equally into validation and test sets (68 kinases in each).

**Fig.4:** Distribution of model predicted LG affinities for the entire train and test set of kinases (left panel) and for one kinase from each group (right panel). Hits and non-hits are shown separately. The bulk of the distribution of predicted affinities for hits is higher than that of non-hits (note the logarithmic scale on the y axis), but many hits have predicted affinities overlapping with the bulk of non-hits. A clean separation only occurs in a subset of cases. We frame the identification of hits based on predicted affinities as a classic binary selection problem, whose outcome depends on the choice of a cutoff. (This plot represents results from run #74839.)

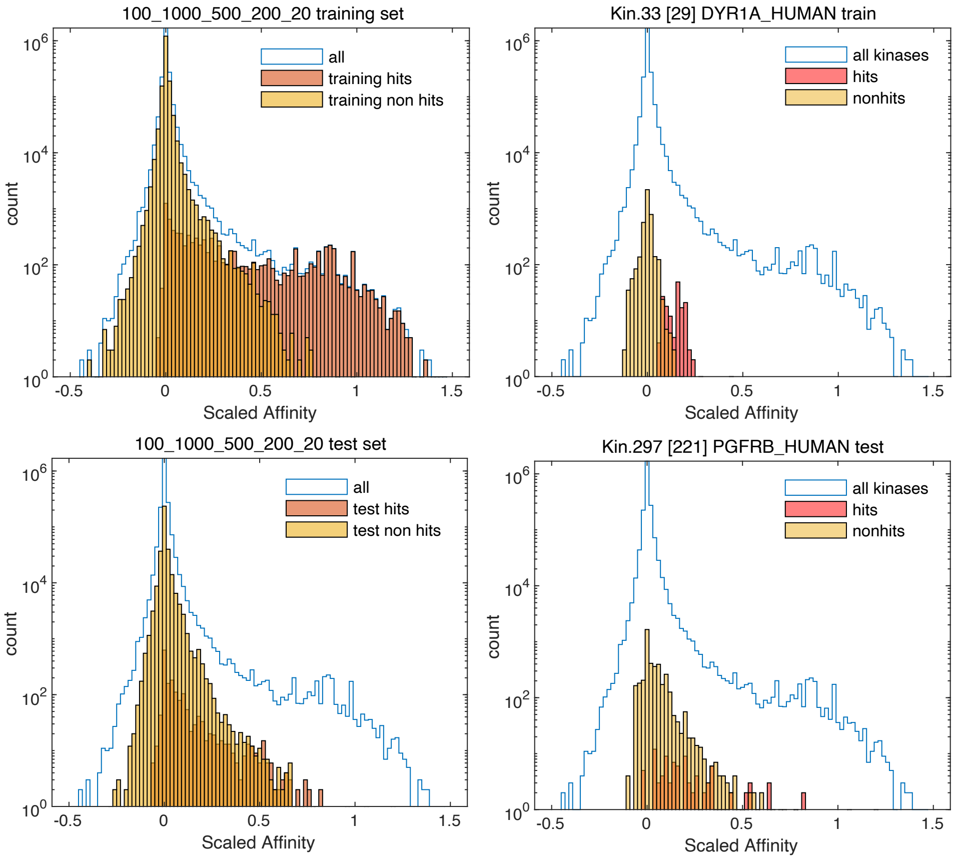

**Fig.5:** Ranges and means of the distribution of (NN) predicted scaled affinities for the hits and nonhits of each of the 455 kinases, from one model run. The ranges typically overlap but the hits are higher than the non-hits. Here, kinases are sorted by the number of known interactions (hits) for the respective kinase, also indicated by the sizes of the solid circle markers. Train, validation, and test groups are indicated by color. (This plot represents results from run #74839.)

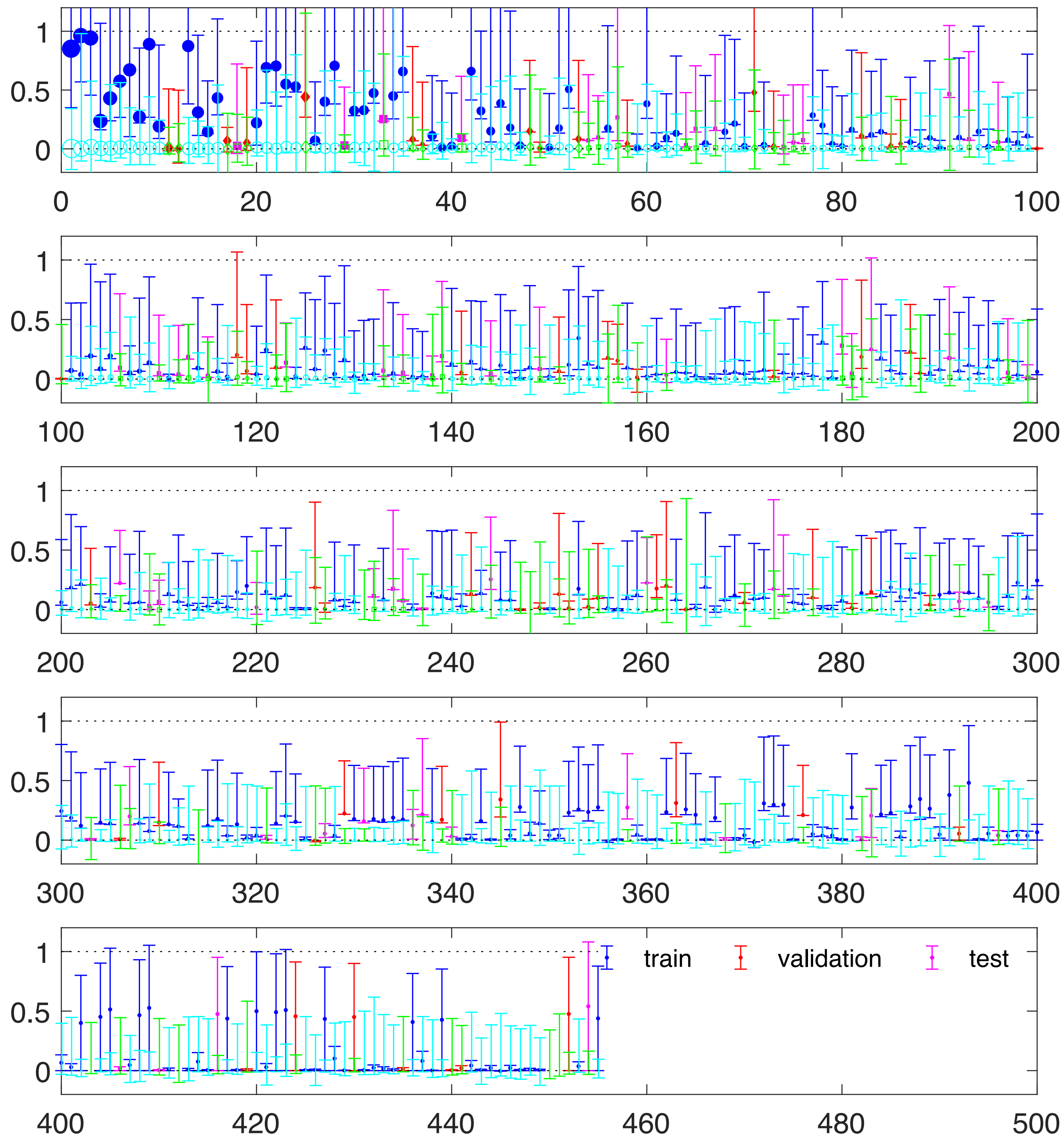

**Fig.6:** Area under the curve (AUC) estimated for individual kinases using the respective receiver operating characteristic (ROC) curves. The left panel shows a scatter plot of the values, with AUC on the y axis and the number of known hits for the respective kinases on the x axis. Train, validation, and test sets are indicated by color. A histogram of the AUC values of the three kinase groups is shown in the right panel. The test set performs worse than the train set (as expected), but the majority of kinases in the test set have AUC above 80-90%, indicating that the predicted affinity values are a good predictor. (This plot represents results from run #74839.)

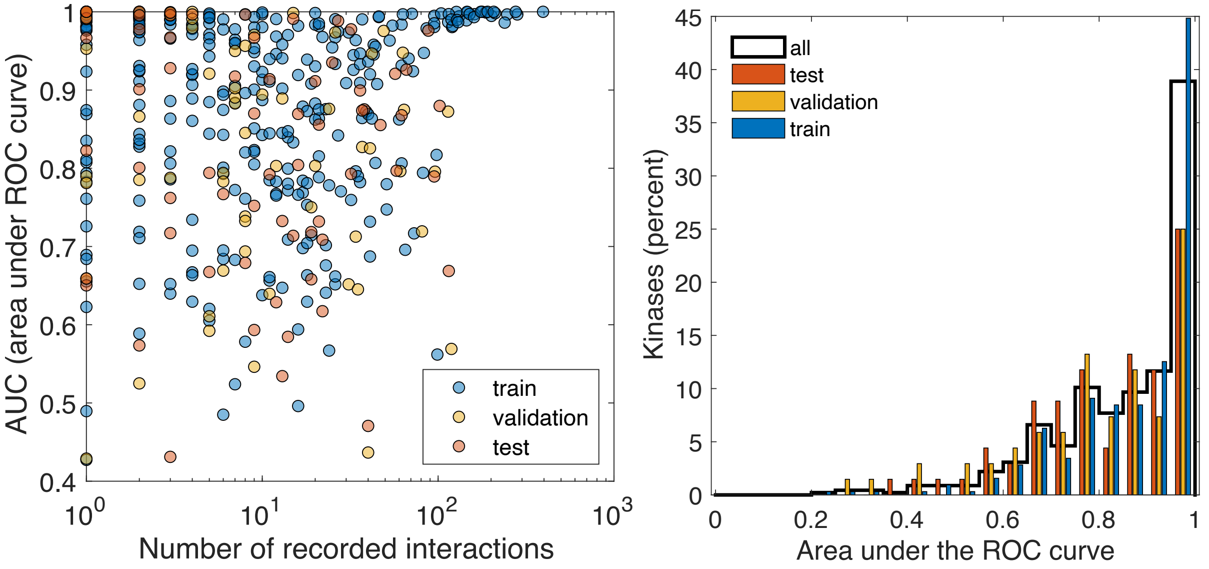

**Fig.7:** Precision and Recall measures obtained using the NN model for one model run (#74839). Precision is calculated for individual (by-kinase) cutoff values that retrieve 50% of the known hits*; Recall is calculated using individual cutoffs at the top 5% of all predicted affinity values* for the respective kinase. Train, validation, and test sets are indicated by color. Marker sizes indicate the number of known interactions ("hits") for the respective kinase. (*Ligand groups that do not interact with the train set kinases are omitted from the hit count of test and validation kinases). Histograms on the sides are marginal distributions and represent the number of kinases with the respective performance measures.

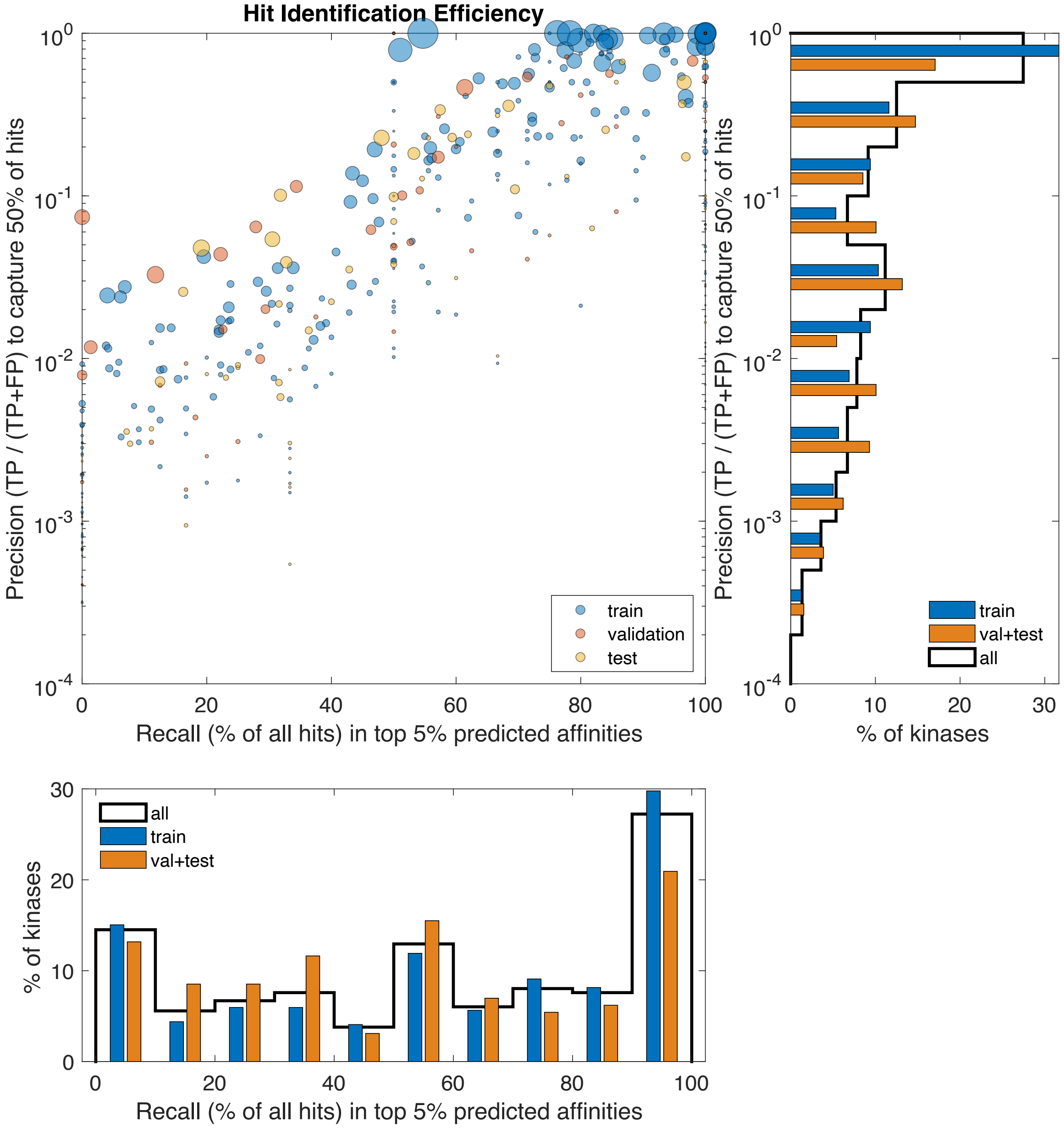

**Fig. 8:** Summary of a performance measure (area under the global ROC curve) for multiple model runs, including NN, linear regression, and combined. All model runs have been performed with the same selection of training (319), validation, and test kinases (68 each); the PCA basis for the output vectors (ligand group affinities) is constructed using the training kinases only, and ligand groups that do not interact with the training kinases are excluded.

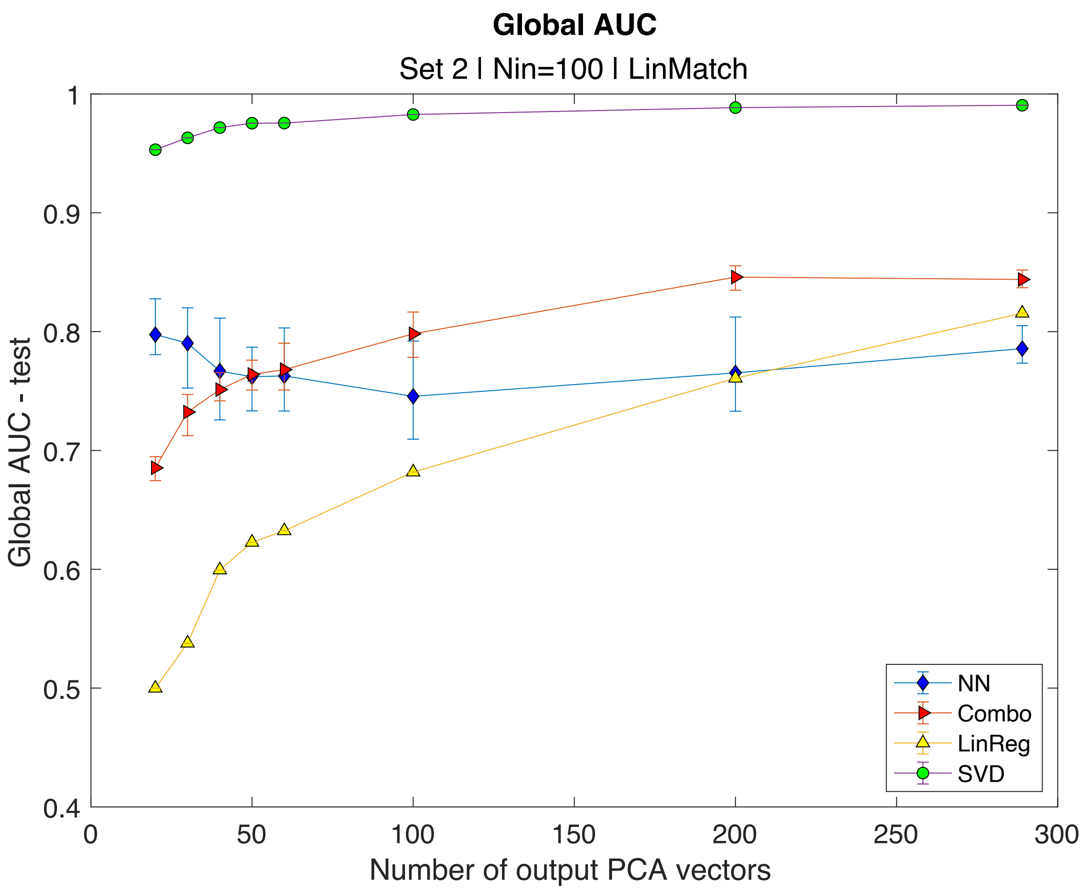

**Table R1:** Values obtained for the AUC, Precision and Recall measures, for one model run (ID 74839, Set 2). Values labelled global were obtained by counting all the hits or non-hits of the respective kinases, above a given cutoff or in total. For the recall, this is equivalent to a weighted average of the by-kinase recall values.

| | AUC (area under the ROC curve) | | |
|---|---|---|---|
| | train | validation | test |
| global | 0.9364 | 0.7787 | 0.8276 |
| mean by kinase | 0.8779 | 0.8068 | 0.8144 |
| mean by hit | 0.9334 | 0.7650 | 0.8190 |

| | Recall | | | |
|---|---|---|---|---|
| Cutoff | global | | mean | |
| | train | test | train | test |
| 1% of all | 0.2859 | 0.2739 | 0.4051 | 0.3689 |
| 5% of all | 0.703 | 0.4897 | 0.5907 | 0.5245 |

| | Precision | | | |
|---|---|---|---|---|
| Cutoff | mean | | median | |
| | train | test | train | test |
| 1% of all | 0.1929 | 0.1284 | 0.0682 | 0.0795 |
| 5% of all | 0.0966 | 0.0468 | 0.0185 | 0.0255 |
| 50% of hits | 0.3326 | 0.1773 | 0.1351 | 0.0434 |

**Table R2:** Comparison of performance measures obtained on the training set for the AUC, Precision and Recall measures, for one NN model run (ID 74839 as described previously); linear regression using all PCA components, and their combination.

| | AUC | | Recall | | Precision | | |
|---|---|---|---|---|---|---|---|
| | global | mean by kinase | Top 1% | Top 5% | Top 1% | Top 5% | 50% of hits |
| NN (100\|20) | 0.8276 | 0.8145 | 0.3689 | 0.5245 | 0.1284 | 0.0468 | 0.1774 |
| LinReg (425\|289) | 0.7960 | 0.8201 | 0.2693 | 0.5112 | 0.1054 | 0.0448 | 0.1392 |
| Combo | 0.8580 | 0.8509 | 0.3234 | 0.5655 | 0.1222 | 0.0535 | 0.1585 |
| SVD (100\|20) | 0.9531 | 0.9336 | 0.4920 | 0.7329 | 0.1780 | 0.0678 | 0.2934 |
